# ionbot: a novel, innovative and sensitive machine learning approach to LC-MS/MS peptide identification

**DOI:** 10.1101/2021.07.02.450686

**Authors:** Sven Degroeve, Ralf Gabriels, Kevin Velghe, Robbin Bouwmeester, Natalia Tichshenko, Lennart Martens

**Affiliations:** VIB-UGent Center for Medical Biotechnology, VIB, Ghent, Belgium; Department of Biomolecular Medicine, Ghent University, Ghent, Belgium

## Abstract

Mass spectrometry-based proteomics generates vast amounts of signal data that require computational interpretation to obtain peptide identifications. Dozens of algorithms for this task exist, but all exploit only part of the acquired data to judge a peptide-to-spectrum match (PSM), ignoring important information such as the observed retention time and fragment ion peak intensity pattern. Moreover, only few identification algorithms allow open modification searches that can substantially increase peptide identifications.

We here therefore introduce ionbot, a novel open modification search engine that is the first to fully merge machine learning with peptide identification. This core innovation brings the ability to include a much larger range of experimental data into PSM scoring, and even to adapt this scoring to the specifics of the data itself. As a result, ionbot substantially increases PSM confidence for open searches, and even enables a further increase in peptide identification rate of up to 12% by also considering lower-ranked, co-eluting matches for a fragmentation spectrum. Moreover, the exclusive use of machine learning for scoring also means that any future improvements to predictive models for peptide behavior will also result in more sensitive and accurate peptide identification.

## Main

Liquid Chromatography combined with high-resolution (Tandem) Mass Spectrometry (LC-MS/MS) has established itself as an invaluable technology for sensitive proteome analysis. It generates vast amounts of raw signal data that require biological interpretation from dedicated bioinformatics tools, known as peptide identification engines. These engines seek to accurately match the observed LC-MS/MS signals with the peptide molecules that generated them. To date, dozens of such identification engines have been developed ^1^.

Nevertheless, peptide identification from LC-MS/MS data remains far from trivial. Spectra are noisy and incomplete, and sequences can be several tens of amino acids long. This leads to an extremely large number of potential target sequences, which in turn poses computational challenges, along with specificity challenges ^2^. As a result, a commonly employed tactic for identification engines is to reduce the search space to only the proteome of interest for the sample under study. Yet, even this strong restriction is typically not sufficient, as several hundreds of potential amino acid modifications should be considered as well, again leading to an enormous growth in the number of possible targets, even for small proteomes. Most of the classical identification engines maintain a more manageable search space by drastically reducing the amount of potential modifications that can be considered. The obvious result is that many relevant modifications may be missed, leading to many false negatives. Recently, several so-called open modification search engines have therefore been developed, that see to address this limitation by allowing the full range of potential modifications to be considered during a fast open modification search that can be applied to large LC-MS/MS datasets ^3–5^.

All identification engines compute a score for each considered peptide-to-spectrum-match (PSM) that is typically designed to reflect the probability of obtaining a true positive identification. As this PSM scoring function decides on the best match for a given MS2 spectrum, the resulting scores should predict the likelihood of a true positive identification as accurately as possible, especially when searching spectra against large search spaces, as is the case for open searches ^2^. With different engines implementing different PSM scoring functions, the set of identified PSMs can be very different between them.

Ideally, the PSM scoring function accurately models the expected LC-MS/MS signal from a given peptide and relies on the comparison between the expected and the observed data to judge a match. This can be achieved in practice by exploiting all accessible PSM data as *matching information*, including the observed retention time for the LC separation, the precursor m/z for the MS1 analysis, and the MS/MS spectrum for the fragmentation analysis. Importantly, Machine learning models already exist to accurately predict calibrated expected retention times ^6–8^ and expected MS2 peak intensities from peptide sequences ^8–11^

However, current identification engines fail to efficiently exploit this matching information in the scoring function. Typically, the PSM score is computed by counting matched peaks, in some way weighted by the MS2 intensities they explain. This implicitly or explicitly penalizes unmatched fragment ions, which is particularly problematic in open searches, where more accurate scoring functions are required. To address this issue and improve accuracy, the relationship between the peptide amino acid sequence and the corresponding peak intensity pattern needs to be considered in the PSM scoring function ^12^.

An ideal PSM scoring function would optimally combine all the relevant sources of matching information into a single accurate score. Substantial progress towards building such a scoring function was made by applying Machine learning to rescore leveraged feature vector representations of PSMs that can contain any source of information ^13,14^. The PSM rescoring function is then computed from these feature vector representations of the experimental data. However, only limited data is explored as only first-ranked PSMs obtained by an engine are rescored. This means that false first-ranked PSMs cannot be replaced by the true PSM based on the leveraged matching information.

We here therefore introduce ionbot, a completely new type of open modification search engine that exploits the ability to incorporate all relevant matching information into a single, Machine learning-based score. To achieve this, we introduce the concept of a candidate match set that is not restricted to first-ranked matches and is computed from a vast open search space using predicted sequence tags ^15^ and a set of biased PSM scoring functions. Furthermore, the ionbot PSM scoring function is computed from this candidate match set using semi-supervised Machine learning, making the PSM scores reproducibly tailored to the experimental data. This leads to an engine that outperforms traditional search engines, allows for reliable open modification searches that outperform current open modification engines, and can be readily adapted to very specific conditions, such as TMT labelled data sets, further increasing identification rates. Finally, we show that our approach naturally leads to the identification of a substantial amount of lower-ranked co-eluting PSMs from chimeric MS2 spectra. Throughout, ionbot maintains a tightly controlled FDR, illustrating superior sensitivity while maintaining specificity.

## Results

### Sequence tag prediction models learn from MS2 peak intensities and show high accuracy

In this section the prefix and suffix tag prediction models (Methods) implemented in ionbot are evaluated on the testing set by the Area under the ROC Curve (AUC) and the Average Precision (AP) computed from the Precision-Recall (PRC) curve. All models show very high predictive accuracy with suffix tag models (AUC=99.9/AP=98.2 for HCD and AUC=99.9/AP=99.4 for HCDTMT) performing better than prefix tag models (AUC=99.8/AP=93.1 for HCD and ACU=99.9/AP=99.2 for HCDTMT) (Supplementary Fig. 1). It is worth noting that TMT trained models show highest predictive accuracy, especially for the prefix tags.

Furthermore, scoring a HCD testing set with a HCDTMT model and *vice versa* substantially decreases predictive accuracy. For the models trained on HCD and evaluated on HCDTMT, the prefix model reduced to AUC=98.9 and AP=58.7, while the suffix model shows a slight decrease to AUC=99.8 and AP=95.6. Notably, for models trained on HCDTMT and evaluated on HCD, the prefix models prediction performance decreased much further to AUC=87.7 and AP=7, with a smaller decrease for the suffix model (AUC=99.5/AP=92.2).

To further evaluate the models, the true PSMs identified by an open search were analyzed. For each true PSM a prefix and suffix tag ranking was computed by scoring all tags with the corresponding predictive models. The highest rank between the true prefix and suffix tag (determined by the matched peptide) is then recorded as a metric for how well the predicted sequence tags can reduce the search space (Methods). The vast majority of these ranks were within the top-10, with many ranked first for one of the two models (Supplementary Fig. 2).

### Expanding the search space is crucial; we recommend to no longer use closed searches

Open searches can match peptidoforms never considered in closed searches. However, at the same time, a larger search space leads to higher scoring decoy matches, thereby potentially increasing the PSM score threshold required to maintain a 1% FDR ^16^. In this section we investigate the difference between a closed and open ionbot search for the five evaluation datasets (Methods).

Our findings confirm previous research showing that open searches considerably increase proteome coverage. The increase in PSM identifications is 52% for HEK239 (Fig. 1a), and up to 70% for TMTCPTAC (Supplementary Fig. 3). At the unique peptide level, identification gains go up to 25% (HEK239) and 27% (TMTCPTAC). It is worth noting that this overall increase in the number of identifications will play an important role in the accurate downstream protein inference and quantification, as well as in increasing the power of the succeeding differential analyses.

**Fig. 1.**
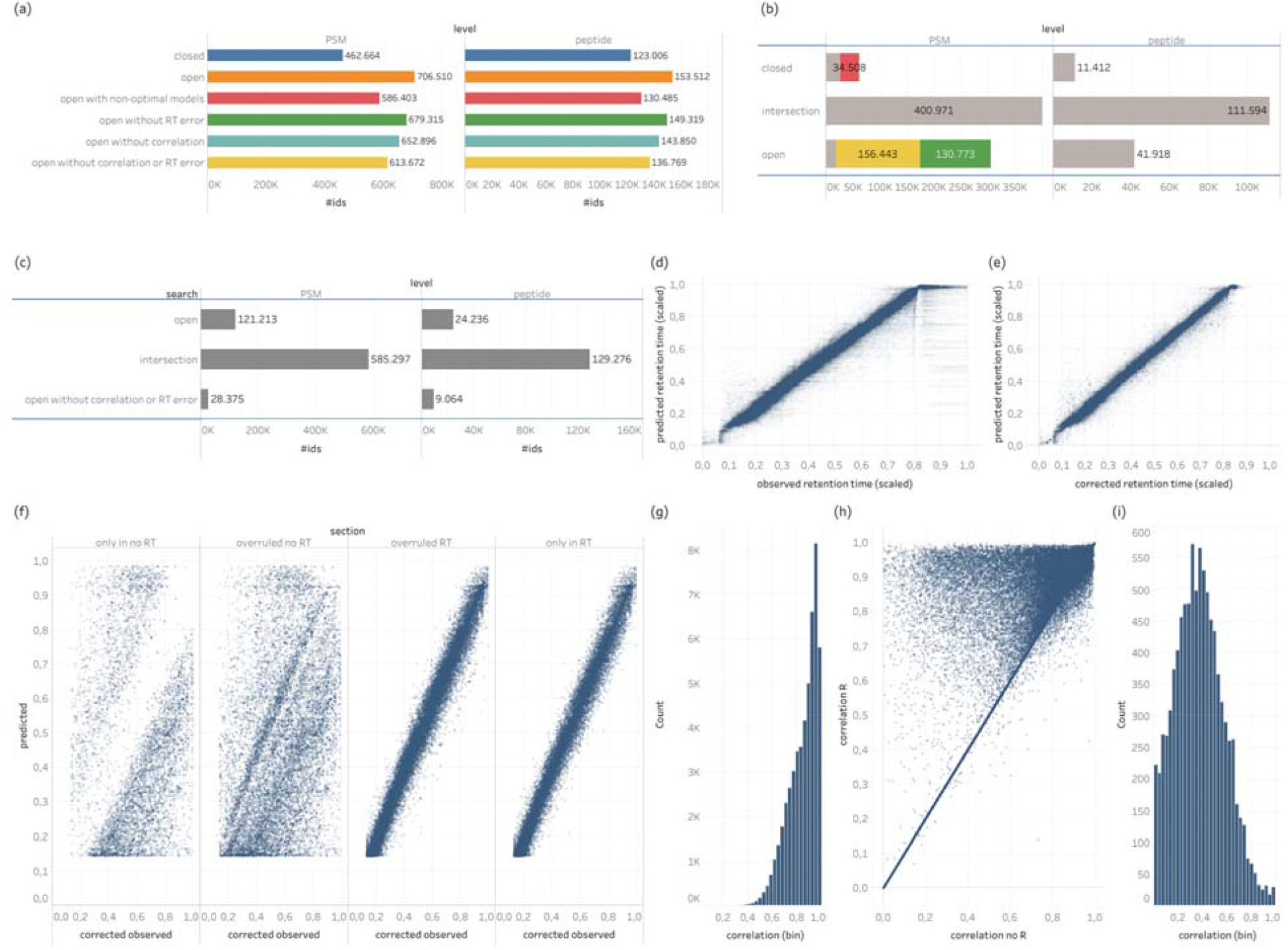
Comparing different implementations of the ionbot search engine. **(a)** shows the number of identifications in HEK239 for the different implementations. These are a closed search (dark blue), a standard open search (orange), an open search using non-optimal models (red) (for the TMTCPTAC dataset, models (tag-models, MS2PIP and DeepLC) trained on unlabeled HCD were used, while for all other datasets models trained on TMT labeled data were used), an open search without using the RT-pred-error feature (green), an open search without using the intensity-correlation feature and biased scoring function (light blue), and an open search without intensity-correlation and RT-pred-error (yellow). **(b)** shows the identification overlap between a closed and open HEK239 search. For the closed searches, spectra with overruled identifications are shown in red. For open searches, matches with an unexpected modification are shown in green, while wide error matches are shown in yellow. **(c)** shows the identification overlap between an open HEK239 search and the same open search without using the intensity-correlation and RT-pred-error information. **(d)** true observed versus DeepLC predicted retention time for CD8T. **(e)** corrected observed versus DeepLC predicted retention time for CD8T. (f) compares corrected observed versus predicted retention times for HEK239 searches using (with-RT) and not using (no-RT) the RT-pred-error feature. **(g)** shows intensity-correlation (R) for PSMs identified in HEK239 identified with using R but not identified not using R (no R). **(h)** plots the intensity-correlations for spectra identified in R and no R, but with a different match. **(i)** shows R for PSMs identified in no R but not identified in R.

Counting PSM and peptide identifications does not reveal all the differences between a closed and open search. We found that a substantial number of closed search identifications are no longer called in the corresponding open search. This is 12% of the closed search identifications for HEK239 (Fig. 1b) and goes up to 19% for the Breast dataset (Supplementary Fig. 5). Furthermore, many of these ‘lost’ matches are overruled by a better match in the open search, as indicated in the figures. It is likely that most of these overruled peptide matches are incorrect and have been forced upon closed search identifications due to the absence of the otherwise higher scoring, true peptide ^17^. This is confirmed by the group FDR computed specifically for overruled PSMs, which is 8.7% for Brain, 10% for HEK239 and CD8T, 12% for Breast, and goes up to 16.6% for TMTCPTAC. Note that PSMs are overruled by a different peptide sequence as we consider PSMs as the same when they match the same peptide sequence with potentially different peptidoforms.

Notably, the majority of PSMs gained in open searches are explained by the wide (7.5 Da) precursor mass error tolerance that ionbot allows for matches without unexpected modification. These precursor errors show a periodic pattern at 1 Da intervals (Supplementary Fig. 6). This is likely due to another peptide within the isolation window that accounts for (most of) the MS2 singals^18^.

### Prediction models trained on specific experimental conditions improve identification

For each predictive model implemented in ionbot (tag-models, MS2PIP and DeepLC), there is a version trained on unlabeled HCD data, and a version trained on TMT labeled HCD data. In this section we apply ionbot with TMT specific prediction models on the non-TMT labeled evaluation datasets. Similarly, we applied ionbot not using TMT-specific models on the TMTCPTAC dataset.

We observed a 17% decrease in PSM and a 15% decrease in peptide identifications when employing non-optimal predictive models in HEK239 (Fig. 1a). For the other datasets the loss in PSMs amounts for 16% for CD8T, 15% for Brain, 24% for Breast, and 14% for TMTCPTAC. At the peptide level these losses are repeated, with a loss of 13% for CD8T, 15% for Brain, 20% for Breast, and 24% for TMTCPTAC (Supplementary Fig. 3-4).

### Predicted retention time and fragment ion intensities provide decisive PSM information

In this section we investigate the relevance of the DeepLC retention time predictions (RT-pred-error) and MS2PIP peak intensity predictions (intensity-correlation) features in the ionbot PSM scoring function (Methods). Grouping PSMs by peptidoform (Methods) to compute the corrected observed retention time to compute RT-pred-error clearly reduced long elution time windows peptidoforms identified by multiple spectra (Fig. 1d-e). This is especially true for peptides at the end of an LC run, where the issue can be even more problematic for the RT-pred-error feature.

Comparing open searches with and without using the RT-pred-error feature in the PSM scoring function showed that consistently more PSMs were identified when the feature is added to the scoring function. At first, this gain appears to be relatively small, at 3.8% for HEK239 (Fig. 1a) and 2.2% for CD8T, 2.8% for Brain, 6.8% for Breast, and 0.6% for TMTCPTAC (Supplementary Fig. 1), but it is worth noting that the vast majority of true PSMs are (also) confirmed by the other sources of matching information, which leaves the retention time feature to correct only ambiguous situations that cannot be distinguished by any of the other sources. We found that PSM identifications unique to the search not using RT-pred-error show high retention time error in general, and, that many PSMs that were overruled when adding the feature show high prediction error as well (Fig. 1f, Supplementary Fig. 8).

Similarly, we compared open searches with and without using the intensity-correlation feature in the scoring function. The latter search also does not use this correlation information as a biased PSM scoring function. We saw an increase in PSMs (7.6%) identified when adding the feature for HEK239 (Fig. 1a). For the other datasets the gain is 5.4% for CD8T, 6.8% for Brain, 6% for Breast, and 10% for TMTCPTAC (Supplementary Fig. 1). Yet here again, this information only gains importance in ambiguous situations that cannot be distinguished by any of the other sources. For HEK239, identifications called only by using the intensity-correlation feature show high correlations (Fig. 1g), while PSMs that are eliminated when the correlation feature is used show low overall correlations (Fig. 1i). Also, for overruled matches when intensity-correlation is used, the difference in correlation can be large, even though the vast majority shows only small differences in the higher correlation range (Fig. 1h). In these cases, it becomes difficult to decide on the correct match based on correlation and other matching information is required to decide on the true match. The same conclusions were made for the other evaluation datasets (Supplementary Fig. 9).

We repeated the same experiment with RT-pred-error and intensity-correlation omitted from the scoring function. This resulted in a much more substantial increase in the number of PSM identifications, with 13% for HEK239 (Fig 1a), 10% for CD8T, 11.8% for Brain, 13.6% for Breast, and 12.4% for TMTCPTAC (Supplementary Fig. 1). At the peptide level the gains amount to 9% for CD8T, 11% for HEK239, 11.6% for Brain, 10.8% for Breast, and 11.5% for TMTCPTAC (Supplementary Fig. 2).

Considering retention time error and intensity correlation in the PSM scoring function not only increases the number of identifications, but also corrects and overrules incorrect matches based on the additional matching information that becomes available. For instance, for HEK239, 2.8% of the PSMs identified omitting both features were overruled by a better match when using them (Fig. 1c). Similar results were found for the other datasets (Supplementary Fig. 10).

Finally, we computed the group FDRs for PSMs that were identified when not using RT-pred-error and intensity-correlation but that were no longer identified when these predictions were added as a biased scoring function and as a component in the PSM scoring function. The group FDRs for these ‘disappearing’ identificationswere again very high, with 6% for Breast, 21% for HEK32, 23.5% CD8T, 24% for Brain, and up to 36% for TMTCPTAC, respectively.

### Identification sensitivity is substantially increased by considering lower rankedco-eluting matches

To determine the best match for an MS2 spectrum, ionbot learns the PSM scoring function from the candidate match set and then selects the first-ranked match for each spectrum based on the computed scores. Next, a more accurate PSM score is computed from these first-ranked matches, and the statistical significance is determined for these first-ranked PSM scores (Methods).

Nevertheless, the candidate match set explicitly contains multiple candidates for many spectra (Methods). In this section we investigate computing the statistical significance of scores from all PSMs in the candidate match set, which can then result in multiple candidate peptides passing the 1% FDR threshold for a given spectrum. We found that even though the FDR threshold does not impose a limit on the number of possible matches, the vast majority of spectra with multiple identified matches had just two (Supplementary Fig. 11). The maximum number of different matches observed for an MS2 spectrum was four.

Considering all co-eluting matches greatly increases identification sensitivity, particularly at the unique peptide sequence level. This gain in unique peptide sequences is 12.4% for Brain, 10.2% for HEK239 (Fig. 2a), 10.3% for CD8T, 2.5% for TMTCPTAC, and 5.1% for Breast (Supplementary Fig. 14).

**Fig. 2.**
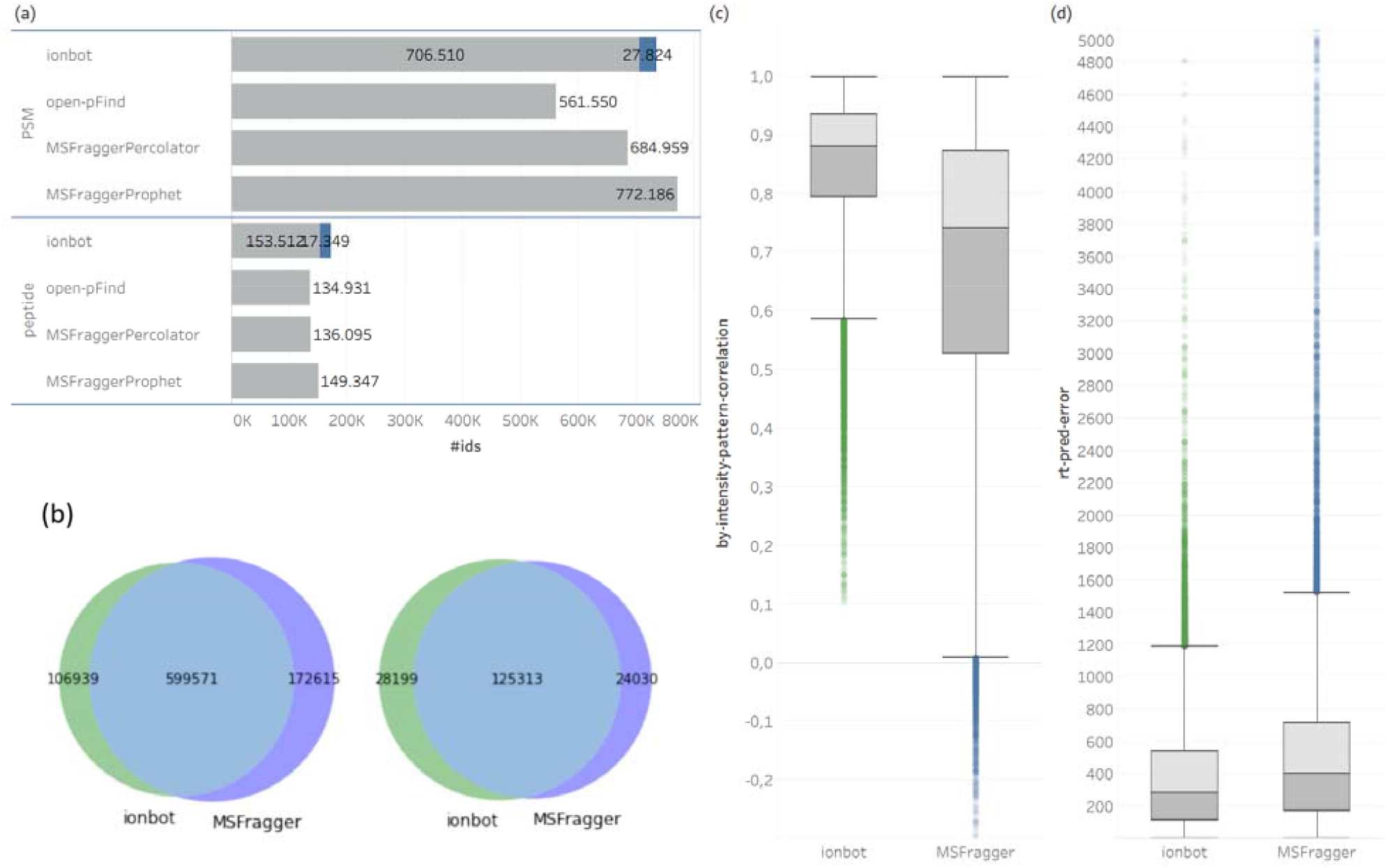
Identification comparison between ionbot and MSFragger for HEK239. **(a)** shows the number of PSM and unique peptide identifications for each engine. **(b)** shows the identification overlap between ionbot and MSFragger at PSM level (left), and at unique peptide sequences level (right). **(c)** shows the value of the intensity-correlation feature for PSMs unique to ionbot or MSFragger. **(d)** shows the value of the retention time prediction error feature for PSMs unique to ionbot or MSFragger.

Despite the fact that similar peptide matches (edit distance < 4) for a given spectrum were filtered out of the candidate match set before learning the PSM scoring function (Methods), there can still be a substantial overlap in fragment ions between co-eluting matches, which leads to the use of the same intensity information more than once. To investigate, we computed the ratio of the intersection of b- and y-ion peak masses over the union of these masses (Supplementary Fig. 12). The results clearly show that the vast majority of these co-eluting matches have a ratio close to zero, indicating almost no overlap between the assigned b- and y-ion peaks.

Universal Spectrum Identifiers ^19^ to spectrum annotations can be found in the Supplementary Tables.

### Entrapment peptides confirm accuracy and stability of the ionbot open search FDR estimates

For the FDR estimates to be meaningful, the ionbot PSM scoring function should treat false matches against the decoy and target database equally, i.e., decoy matches should be representative random matches. For ionbot, this conveys that the PSM scoring function learned from experimental data should not be biased towards favoring matches against the target database.

To estimate a potential matching bias, we adopted the entrapment peptides approach that adds sequences of evolutionary sufficiently different species to the target database to capture random matches against the true target database^20^ (Methods). If a bias exists, we should observe more than the expected number of matches against the entrapment compared to the decoy database. For the *Pyrococcus furiosus* sample, the entrapment database (279.618 sequences) is significantly larger than the true target database (2045 sequences), so we expect to see about the same ratio of entrapment as of decoy hits for random matches. Note that peptides shared between true and entrapment database are counted as matches against the true database. The human and mouse target databases both contain a similar number of true target, and of *Archaea* entrapment sequences. In this case we expect to see about twice as many decoy hits compared to entrapment hits for random matches. Supplementary Fig. 13 plots the PSM q-values against the entrapment FDR for each of the six datasets.

When we compare the first ranked matches for ionbot and MSFraggerPercolator we observe very similar results for all datasets. For the five evaluation datasets we observe, at q-value = 1%, that first ranked ionbot matches show an entrapment FDR of 0.43% for Breast, 0.44% for CD8T, 0.47% for TMTCPTAC, 0.5% for Brain, and 0.51% for HEK239. This even goes as low as 0.23% for the *Pyrococcus furiosus* dataset.

For MSFraggerProphet no q-values were available. We observe surprisingly small entrapment FDRs for Brain (0.02%), HEK239 (0.03%), and TMTCPTAC (0.03%). However, for the *Pyrococcus furiosus* dataset, the entrapment FDR is much higher than expected at 2.4%.

With the exception of the Brain dataset, the lower ranked ionbot matches show slightly higher entrapment FDRs compared to the first ranked matches. For the five evaluation datasets we observe, at q-value = 1%, an entrapment FDR for lower ranked ionbot matches of 0.48% for Brain, 0.58% for TMTCPTAC, 0.65% for CD8T, 0.84% for HEK239, and 1.4% for Breast. For the *Pyrococcus furiosus* dataset this goes up to 2.4%. However, note that for this dataset, there were only 257 lower ranked co-eluting matches identified with just 7 PSMs matching the entrapment database. It is therefore very difficult to obtain an accurate estimate.

### The ionbot engine compares favorably to other state-of-the-art open modification engines

Many open search engines exist, but few can produce sensitive identification results for large datasets that contain hundreds of thousands of spectra, mainly due to computational limitations. Two recent engines stand out in terms of identification sensitivity and speed: MSFragger and open-pFind (Methods). In terms of speed, MSFragger is well known for its fast implementation of the open search. On a 24 CPU machine, MSFragger takes 9 minutes to process a 50k spectra dataset in open search. It took ionbot 24 minutes to process that same dataset with the same settings. However, even though ionbot is 2.5 times slower than MSFragger, ionbot is still sufficiently fast to process large dataset within acceptable time.

We first examine first ranked matches only. Note that the number of identifications obtained with PeptideProphet or Percolator can be very different, with an 11% difference for the HEK239 dataset (Fig. 2a and Supplementary Fig. 14). At the PSM level, both ionbot and MSFraggerProphet show substantially higher identification rates compared to open-pFind, mainly due to the 7.5 Da wide error matches (data not shown) that are considered only in ionbot and MSFragger. At the peptide level, ionbot identifies substantially more unique peptides compared to the other search engines (except for the Breast dataset), in turn resulting in more protein identifications (Supplementary Fig. 18).

Next, we focused on comparing ionbot and MSFraggerProphet.We plotted the PSM and peptide identification overlap, which reveals a notable level of disagreement between these two search engines (Fig. 2b, Supplementary Fig. 15). To obtain more insight into this disagreement, we looked at the intensity-correlation computed for PSMs uniquely identified by one of the engines (Supplementary Fig. 16). To avoid bias because of unknown effects of specific modifications on the peak intensity pattern, we limited this investigation to identifications without an unexpected modification. We found that many identifications unique to MSFragger are questionable. For HEK239 (Fig. 2c), when we look at the PSMs unique to ionbot, the 25% with the lowest intensity-correlation still show correlation within [0.59,0.9] (excluding outliers). For the PSMs unique to MSFragger this interval falls substantially lower at [0.01,0.53].. The same observations were made for the other datasets, with the 25% unique matches with the lowest peak intensity correlation for MSFragger falling within the interval [0.07,0.57] for CD8T, [0.3,0.65] for Brain, [0.13,0.54] for Breast, and [0.36,0.68] for TMTCPTAC. For the RT-pred-error information we observe less pronounced differences between the search engines (Fig 2d and Supplementary Fig. 17).

Finally, the number of protein group identifications is plotted in Supplementary Fig. 18. For the five evaluation datasets we observe an increase in the number of protein groups identified when using all identified PSMs (first and lower ranked) during protein inference. Comparing protein group identifications with MSFragger results in more protein groups for MSFragger in the HEK239, Brain and Breast dataset, but less in CD8T and TMTCPTAC. It should be noted here that the protein inference algorithms also differ between the two engines, and that this may also have an effect on the final result.

## Discussion and conclusion

We presented ionbot, a novel type of open identification engine that is unique in taking full advantage of the accurate predictions provided by Machine learning algorithms. This is realized by omitting the predefined PSM scoring function altogether, and instead relying on a data-driven strategy to learn the weights of all matching information provided which includes the MS2 peak intensity pattern correlation and LC retention time prediction error. This provides ionbot with the flexibility to adapt the PSM scoring function to the experimental conditions and passes the identification bias from the expert to the data.

We have shown that ionbot exploits the additional matching information in its PSM scoring function to increase the number of identifications at the PSM as well as at the peptide level, and that it also allows ionbot to adapt to specific experimental protocols such as TMT-labelling with ease. Moreover, we show that the additional matching information is also highly useful in reducing false positive matches by either eliminating these, or by replacing them with higher scoring identifications.

When compared to other open modification search engines, ionbot performs on par at the identification sensitivity level. However, looking at the PSM evidence in terms of matching information reveals that many identifications unique to the other engines show very low intensity pattern correlation. This can indeed be expected as other open search engines do not exploit these sources of matching information in their scoring function and, as such, are unable to make accurate decisions based on these sources.

Finally, co-eluting lower-ranked PSMs arise naturally from the data-driven identification strategy implemented in ionbot. The results of the entrapment peptide experiments show that ionbot correctly estimates the FDR for both first ranked matches and lower ranked co-eluting matches. By considering all these co-eluting matches, ionbot can provide substantially higher identification sensitivity compared to the other open search engines.

We believe this research opens up new avenues for improving the analysis of LC-MS/MS data. The Machine learning models employed by ionbot can likely still be improved, especially in terms of predicting the effects on analyte behavior of different protein modifications. Moreover, different (non-)linear models can be evaluated for learning the PSM scoring function. Because ionbot can very easily be fitted with new or improved models, and because ionbot is implicitly highly adaptive, any such improvements will very likely result in more accurate PSM scoring functions. Finally, this highly adaptive nature of ionbot can also allow dedicated optimization for specific experimental conditions, thereby increasing identification sensitivity and accuracy even further.

## Methods

### The search space

Observed MS2 spectra are matched against a peptide search space that is computed in silico from a known proteome using a specific simplified enzymatic cleavage pattern (e.g., for trypsin). In the Human proteome dataset used in this research (see section Datasets), cleaving after lysine or arginine results in 512.990 unique peptide sequences. This number increases to 2.148.714 by allowing for two missed cleavages.

Furthermore, proteins can undergo chemical modifications, with some modifications observed more frequently (e.g. oxidation) than others ^21^. Public repositories that list previously observed modifications, such as unimod.org, currently contain more than 1000 different modifications that are known to alter a specific amino acid in a peptide. Therefore, the peptide search space should be expanded by all possible modified versions of each peptide. Each modification changes the m/z pattern of the MS2 spectrum generated by the modified peptide and can, potentially, also alters the MS2 peak intensity pattern ^22^ and expected LC retention time ^6,23^. To make matters worse, candidate matches for an MS2 spectrum can become very hard to distinguish due to the expanded search space ^2^

The vast majority of peptide identification engines consider only a few of the most expected protein modifications. This is known as a *closed search*. In contrast, in an *open search* one tries to consider all possible peptidoforms (considering all possible modification patterns for a peptide) by adding these to the search space. Even though this space is still considerably smaller than the one evaluated by *de-novo* identification engines (which consider all possible peptide sequences as well), as the number of modifications taken into account increases, the search space expands exponentially, putting a strong computational load on the peptide identification task.

To reduce this computational burden, ionbot limits the open search by setting a maximum on the number of modifications allowed within one peptide. In a closed search, the space consists of all peptidoforms that carry at most two expected (variable) modifications simultaneously. Next, to an unlimited number of expected modifications that are fixed. In an open search, the space is further expanded by all peptidoforms that can be constructed by (i) adding one of the 1000+ post-translational, chemical derivative, or artefactual peptide modifications listed in the unimod.org repository, (ii) considering the delta-masses generated by the modifications in (i) only at the MS1 level, (iii) adding a single amino acid substitution, or (iv) by removing any number of N-terminal amino acids from the peptide to obtain a semi-tryptic peptide. For (i) and (iii), all possible unmodified locations for the modification or substitution are considered. For (ii) ionbot tries to match delta-masses observed only at the MS1 level and therefore not part of the MS2 fragmentation spectrum. For (iv), semi-tryptic peptides are considered up to a length of seven amino acids.

### The candidate match set

The PSM scoring function in ionbot is learned from a candidate match set that is a small subset of the search space, while still being large enough to still contain the true PSMs and to learn an accurate scoring function. To achieve this, ionbot employs a sequence tag strategy driven by Machine learning scoring models, and a set of biased expert-driven PSM scoring functions.

A sequence tag is a short amino acid sequence with a prefix mass and a suffix mass that allocates its position within a peptide. Tags are typically computed from a graph representation of an MS2 spectrum ^24,25^. This spectral graph exploits fragment ion mass differences but ignores the relationship between the tag’s amino acid sequence and the observed MS2 peak intensity pattern. Intensity information is typically exploited only to prefer tags that match higher-intensity peaks.

Instead, ionbot implements a data-driven approach to construct constrained sequence tags. By constraining the tags to the first three (prefix) or the last four (suffix) amino acids in a tryptic peptide, predictive Machine learning models can accurately score a tag from a feature vector representation of the tag and the corresponding MS2 spectrum, thereby eliminating time consuming spectral graph construction, while also exploiting MS2 peak intensity information. For each observed MS2 spectrum, the prefix model scores 8000 3-mers and the suffix model scores 16.000 4-mers, all constructed from nineteen amino acids – structural isomers leucine and isoleucine are treated as a single residue – and oxidized methionine, which was added due to its high prevalence. For each MS2 spectrum, the *T*-top scoring tags from both the prefix and the suffix model are used to reduce the search space to those peptides that match any of these tags within a user-defined MS1 mass error tolerance. For candidate peptides with only fixed, expected or no modifications, this mass error tolerance is increased to 7.5 Da, as justified in the Results section. The parameter *T* should be chosen large enough such that only unlikely peptides are removed from the search space. Here, we set *T*=50 as increasing *T* resulted in just a small identification increase (1%-3%) for all datasets while substantially increasing compute time.

The implementation of a set of biased PSM scoring functions further differentiate ionbot from traditional identification engines. These biased functions each consider just one source of matching information and are listed in Table 2. For peak counting and explained intensity information, ionbot also considers different subsets of fragment ion types. The candidate match set is then further reduced to the top-ranked match for each biased PSM function (when there are multiple matches with the same top-ranked score then all these are kept in the candidate match set).

**Table 1.**
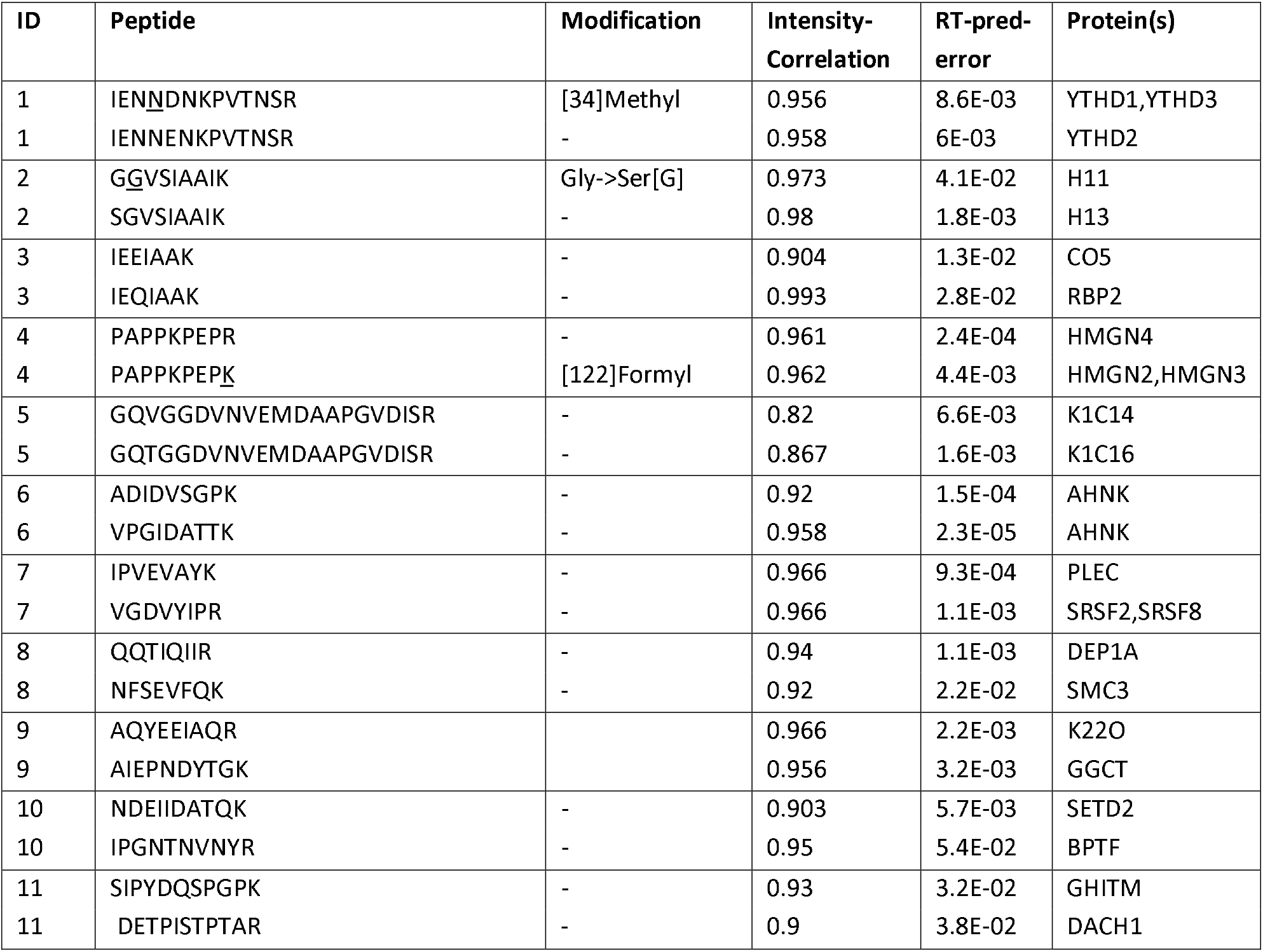
Examples of MS2 spectra with two co-eluting matches. Every ID in the table shows two matches for one MS2 spectrum. For each match, the table shows the peptide sequence (Peptide), its modifications to form the identified peptidoform (Modification), the intensity-correlation, the RT-pred-error (divided by the maximum retention time observed in the corresponding dataset), and the protein(s) that contain the peptide. IDs 1—5 are examples of highly similar matches. IDs 6—11 show matches with highly dissimilar matches.

**Table 2.** Biased PSM scoring functions that filter the search space. Each function considers some plausible form of matching information.

| Biased scoring function | Description |
| --- | --- |
| b1y1-peak-count | total number of b- and y-ions matched considering only charge 1+ |
| b1b2y1y2-peak-count | total number of b- and y-ions matched considering charge 1+ and 2+ |
| peak-count | total number of X-ions matched with X in [a,b,b-H2O,b-NH3,c,x,y,y-H2O,z], considering charge 1+ and 2+ |
| b1y1-intensity-explained | total explained intensity of matched b- and y-ions matched considering only charge 1+ |
| b1b2y1y2-intensity-explained | total explained intensity of matched b- and y-ions matched considering charge 1+ and 2+ |
| intensity-explained | total explained intensity of matched X-ions matched with X in [a,b,b-H2O,b-NH3,c,x,y,y-H2O,z], considering charge 1+ and 2+ |
| intensity-correlation | Pearson correlation between observed and MS2PIP predicted b- and y-ion intensities |
| average-rank | ranks the PSMs by the average rank for ions-count, ions-explained and intensity-correlation |

We observed that in many cases, there can be multiple candidates with highly similar peptides matched for a spectrum. Some examples are shown in Table 1 (IDs 1-5). We filter these out of the candidate match set before learning the PSM scoring function using the following rule: ifa given spectrum matches multiple peptide with edit (Levenshtein) distance three or lower, then keep the peptide match that did not require an unexpected modification. For example, in Table 1 for ID 1, the methylated peptide is removed from the candidate match set. All similar matches without unexpected modification are maintained. For example, for IDs 3 and 5 in Table 1, the candidate match set contains both matches.

### The prefix and suffix tag prediction models

This section explains how the prefix and suffix models score the corresponding 3-mer and 4-mer tags. Both are two-class classification models trained on publicly available spectral libraries (see section Datasets). For a given MS2 spectrum and tag, each model computes the probability that the *k*-mer matches the peptide that generated the spectrum. For the prefix model this is the 3-mer corresponding to the N-terminal three amino acids in the peptide, and for the suffix model this is the 4-mer corresponding to the C-terminal four amino acids in the peptide. The suffix model is designed specifically for peptides that are tryptic at the C-terminal. Even though the prefix model can find tags for peptides not tryptic at the N-terminal, it is important to note that we designed the current version of ionbot for identifying tryptic peptides. The feature vector representation implemented in these models contains discriminative information about both the spectrum and the tag. This allows for modelling the relationship between the tag’s amin acid sequence and the observed MS2 peak intensities.

To represent the tag, a one-hot-encoding of the *k*-mer amino acids was considered, but this yielded much inferior results. Instead, as implemented in our MS^2^PIP tool ^11^, amino acids in the *k*-mer are encoded by each of five amino acid properties: mass, hydrophobicity, helicity, basicity and iso-electric point estimates.

The MS2 spectrum is represented by the observed peak intensities, normalized by total ion current of the spectrum ^26^, of the relevant fragment ions that can be explained by the tag. These are the first three a-, b-, and c-ions for the prefix model, and the first four x-, y- and z-ions for the suffix model, both with variable H2O- and NH3-losses, and with fragment ion charge states 1+ and 2+ considered as well. Variable methionine oxidation is also encoded by adding three binary features that indicate the presence of an oxidation at any position in the tag (excluding the last amino acid in the suffix model tag, which is restricted to lysine or arginine). Lastly, the precursor charge state and precursor mass are encoded as features as well.

Each PSM in the training set is represented by a feature vector for each model, which are collectively labeled as the positive class. For the negative class, the MS2 spectrum is matched to *N* different random *k*-mers from which features vectors are created. For the suffix model, these negative examples remain constrained to lysine or arginine at position four. We found *N*=200 to be a sufficient number of negative *k*-mer examples, while balancing with the computational cost of computing the models.

As in MS^2^PIP, accurate predictive gradient boosted tree models (GBT) are learned with the open-source XGBoost tool ^27^. The boosting algorithm fits an additive decision tree ensemble on the data that allows for extremely fast predictions, which is crucial, considering that ionbot computes hundreds of millions of predictions for a typical spectrum file.

To train and evaluate the prefix and suffix tag scoring models (Methods), the Orbitrap-HCD-best and Orbitrap-HCD-TMT-10 datasets (Table 4) were split into a training and testing (5%) set. The split was computed such that both the positive and all negative feature vectors computed from a tag are either in the training or in the testing set. GBT hyperparameters were optimized using cross-validation on the training set.

**Table 3.** The 43 features used in the ionbot PSM scoring function feature vector. Note that XY-ion-count and XY-ion-explained each constitute 18 different features.

| Name | Description |
| --- | --- |
| charge | charge state of match |
| peplen | length of matched peptide |
| precursor_mass | MS1 precursor mass |
| num_peaks | number of peaks in the MS2 spectrum |
| max_peak | intensity of highest peak in MS2 spectrum |
| XY-ion-count | number of XY-ions matched with X in [a,b,b-H2O,b-NH3,c,x,y,y-H2O,z] and Y charge state +1 or +2 |
| XY-ion-explained | total explained intensity for the XY-ions matched with X in [a,b,b-H2O,b-NH3,c,x,y,y-H2O,z] and Y charge state +1 or +2 |
| intensity-correlation | Pearson correlation between observed and MS2PIP predicted b- and y-ion intensities |
| RT-pred-error | difference between corrected observed and DeepLC predicted retention time |

**Table 4:** Spectral libraries used to train the prefix and suffix models. For each library the number of computed feature vectors labeled as positive (#Positives) and computed feature vectors labeled as negative (#Negative) is show. Libraries were downloaded from chemdata.nist.gov.

| Dataset | Version | #Positives | #Negatives |
| --- | --- | --- | --- |
| Orbitrap-HCD-best | 05-19-2020 | 212.761 | 42.531.154 |
| Orbitrap-HCD TMT-10 | 10-30-2019 | 318.972 | 63.794.400 |

### The PSM scoring function

Ionbot adopts the semi-supervised learning approach pioneered by the Percolator tool ^28^. Each PSM in the candidate match set is encoded as a 43-dimensional feature vector in which each feature represents a different source of matching information. These features are listed in Table 3. The 36 X-ions-counts and X-ions-explained features each consider one type of fragment ion. Combining these into one PSMs scoring function together with the other features is left to the semi-supervised the Linear Support Vector Machine (LSVM) algorithm implemented in Percolator. For the intensity-correlation feature, MS2PIP is called to predict all b- and y-ion peak intensities with the Pearson correlation between the predicted and observed intensities as the value. The RT-pred-error feature is computed as the difference between the corrected observed retention time and the calibrated prediction of DeepLC. The correction is obtained by grouping the PSMs by peptidoform and then, for each group, using the retention time of the PSM with the lowest prediction error as the corrected observed retention time. As shown in the Results section, this substantially increases the relevance of the LC error feature in the PSM scoring function as it is more robust towards peptidoform with long elution windows.

The RT-pred-error feature requires calibration to correct for different experimental conditions. This calls for a limited set of highly confident true PSMs obtained without using the feature. This is realized by first learning the PSM function without the RT-pred-error feature, then selecting the 1000 highest scoring first-ranked matches to calibrate the DeepLC predictions and add the feature to the scoring function. This function is then learned again from the candidate match set.

Finally, ionbot follows the well-established concatenated target-decoy method for computing significance statistics to call true PSMs at a controlled FDR. This means that the learned PSM scoring function selects the highest scoring PSM for each spectrum, from which q-values and other statistics are computed.

Note that ionbot is not restricted to using the Percolator tool. Other Machine learning algorithms can be applied to learn the PSM scoring function as well, including unsupervised and/or non-linear models. Here, we limited ionbot to the LSVM as it is already well-established in the Proteomics community.

### Entrapment peptide database

The entrapment peptides FDR estimation evaluation procedure was adopted from ^20^. Herein, target sequences from evolutionary sufficiently different species are added to the target database, known as the entrapment peptides, that act as true target peptide. The *Pyrococcus furiosus* sample was downloaded from PRIDE (PXD001077) and contains 15.365 MS2 spectra. This spectrum file was searched against a target database that consists of 2045 *Pyrococcus furiosus* proteins and 339 know contaminant proteins as the expected proteins (UniProt reviewed), extended with 193.521 eukaryote and 86.097 vertebrate proteins as entrapment sequences.

The entrapment peptides evaluation procedure was also implemented for the five evaluation datasets in this manuscript by adding 19.678 *Archaea* proteins to the human and mouse target databases. The decoy databases were constructed by shuffling the sequences in the extended target databases.

The result plots (Supplementary Fig. 13) were constructed by sorting the obtained PSMs by q-value (x-axis) and then computing the entrapment FDR

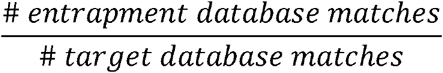

that corresponds to each q-value.

### MSFragger and open-pFind search settings

We compared ionbot with MSFragger (version 3.1 in fragpipe 14.0) and open-pFind (pFind version 3.1.5). We set the following settings equal for each search engine: 20ppm precursor mass error tolerance, peptide length in [7,30], delta-mass (mass range allowed for unexpected modifications) in [-150Da,500Da], a maximum of two missed cleavages, oxidation of M as expected and carbamidomethylation of C as fixed modification. Other settings were left to the default open search settings for each engine.

For MSFragger, we evaluated two post-processing methods: the default PeptideProphet method, and the Percolator method. For the latter we employed Percolator on the .pin files written by MSFragger for further post-processing. The full command line command used to run both versions of MSFragger is included as Supplementary Notes.

## Datasets

### Spectral libraries

The Human Orbitrap-HCD-best and Orbitrap-HCD-TMT-10 spectral libraries were downloaded from the NIST Libraries of Peptide Tandem Mass Spectra^1^. PSMs in these libraries were filtered to contain unique peptide sequences that end with a lysine or arginine only. For peptides that are matched with different spectra, one is selected at random to avoid peptide bias. PSMs are encoded as feature vectors as described in the Methods section. Table 4 shows the number of positive and negative examples in each dataset for training the prefix and suffix tag models.

### Evaluation projects

To evaluate ionbot, we selected five MS2 HCD spectrum datasets of different sizes that were obtained from different labs and with different overall experimental conditions (Table 5). One dataset (Breast) was labeled with super-SILAC and another (TMTCPTAC) with TMT10. All RAW files were downloaded from the PRIDE repository ^29^ and converted to the Mascot MGF format using ThermoRawFileParser ^30^ with the MS2 peak picking option enabled. Spectrum files belonging to the same sample were merged.

**Table 5:** Evaluation datasets used in the Results section. The first four datasets were downloaded from the PRIDE repository (Accession). The TMT datasets was downloaded from the CPTAC portal.

| Name | Accession | #MS/MS spectra | Sample | Machine |
| --- | --- | --- | --- | --- |
| HEK293 | PXD001468 | 1.121.149 | HEK293 cell | Orbitrap Q-Exactive |
| CD8T | PXD000561 | 406.913 | CD8T cell | Orbitrap Elite |
| Brain | PXD001250 | 669.350 | Mouse brain | Orbitrap Q-Exactive |
| Breast | PXD000815 | 382.022 | Breast cancer tumor | Orbitrap Q-Exactive |
| TMTCPTAC | PXD012703 | 1.064.359 | Tumor tissue | Orbitrap Q-Exactive Plus |

## Footnotes

1 https://chemdata.nist.gov

