## Supplementary material for "ionbot: a novel, innovative and sensitive machine learning approach to LC-MS/MS peptide identification": Suppl. Notes

Command line commands used to execute MSFragger:

$ philosopher workspace --clean

$ philosopher workspace --init

$ philosopher database --custom db.fasta --prefix decoy_

Let db_fragger.fas be the created database file. Add it to open_fragger.params and CrystalC.params.

$ java -Xmx500g -jar MSFragger-3.3.jar open_fragger.params *.mzML

This writes .pepXML, .pin and .tsv result files in the folder that contains the .mzML files. To run Crystal-C copy the jar and parameter file to the folder that contains the .pepXML files.

$ java -Xmx500G -cp "CrystalC-1.4.2.jar" crystalc.Run CrystalC.params *.pepXML

This creates *_c.pepXML files.

$ python3 ionbot/scripts/merge.py *.pin > pin

$ percolator --unitnorm --only-psms --post-processing-tdc pin > outfile

Note that these matches still need to be corrected for by Crystal-C, this happens in the notebook and uses the matches obtained with PeptideProphet.

Back in the tools/MSFragger-3.3/ folder:

$ philosopher peptideprophet --nonparam --expectscore --decoyprobs --masswidth 1000.0 --clevel -2 --decoy "decoy_" --combine --database db_fragger.fas *_c.pepXML

This writes interact.pep.xml in the folder that contains the _c.pepXML files.

$ philosopher proteinprophet --maxppmdiff 2000000 --output combined path/interact.pep.xml

This writes combined.prot.xml in tools/MSFragger-3.3/

$ philosopher filter --sequential --razor --mapmods --tag decoy_ --pepxml path/interact.pep.xml --protxml ./combined.prot.xml

or

$ philosopher filter --mapmods --tag decoy_ --pepxml path/interact.pep.xml

Finally write tsv result files in tools/MSFragger-3.3/

$ philosopher report
