## Supplementary material for "ionbot: a novel, innovative and sensitive machine learning approach to LC-MS/MS peptide identification": Suppl. Figures

Supplementary Figures

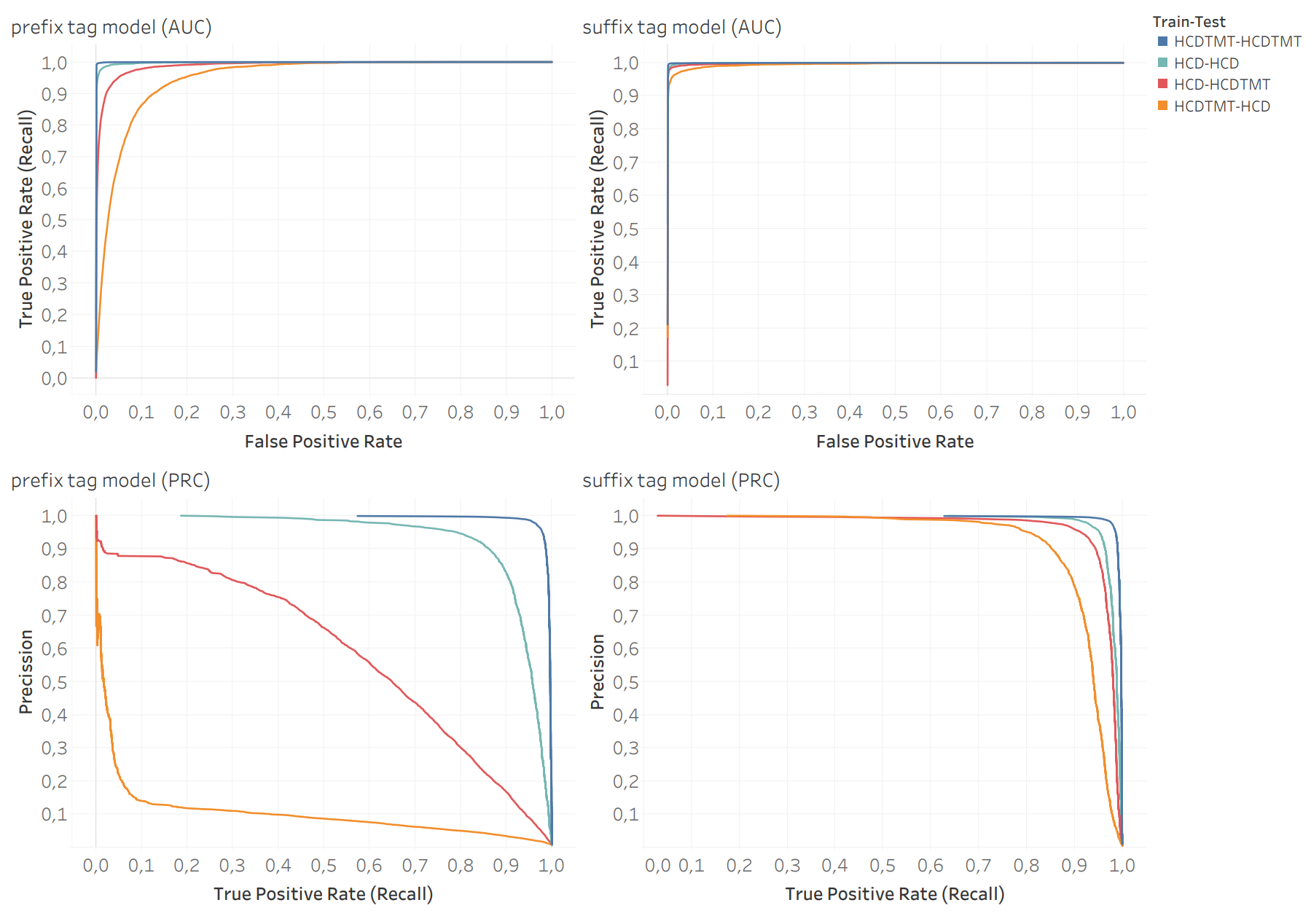

**Supplementary Fig. 1 | AUC and PRC curves for the HCD and HCDTMT prefix and suffix tag scoring models.** The legend explains what training and testing set was used (e.g. for HCD-HCDTMT, the GBT model was fitted on the HCD training set and evaluated on the HCDTMT testing set).

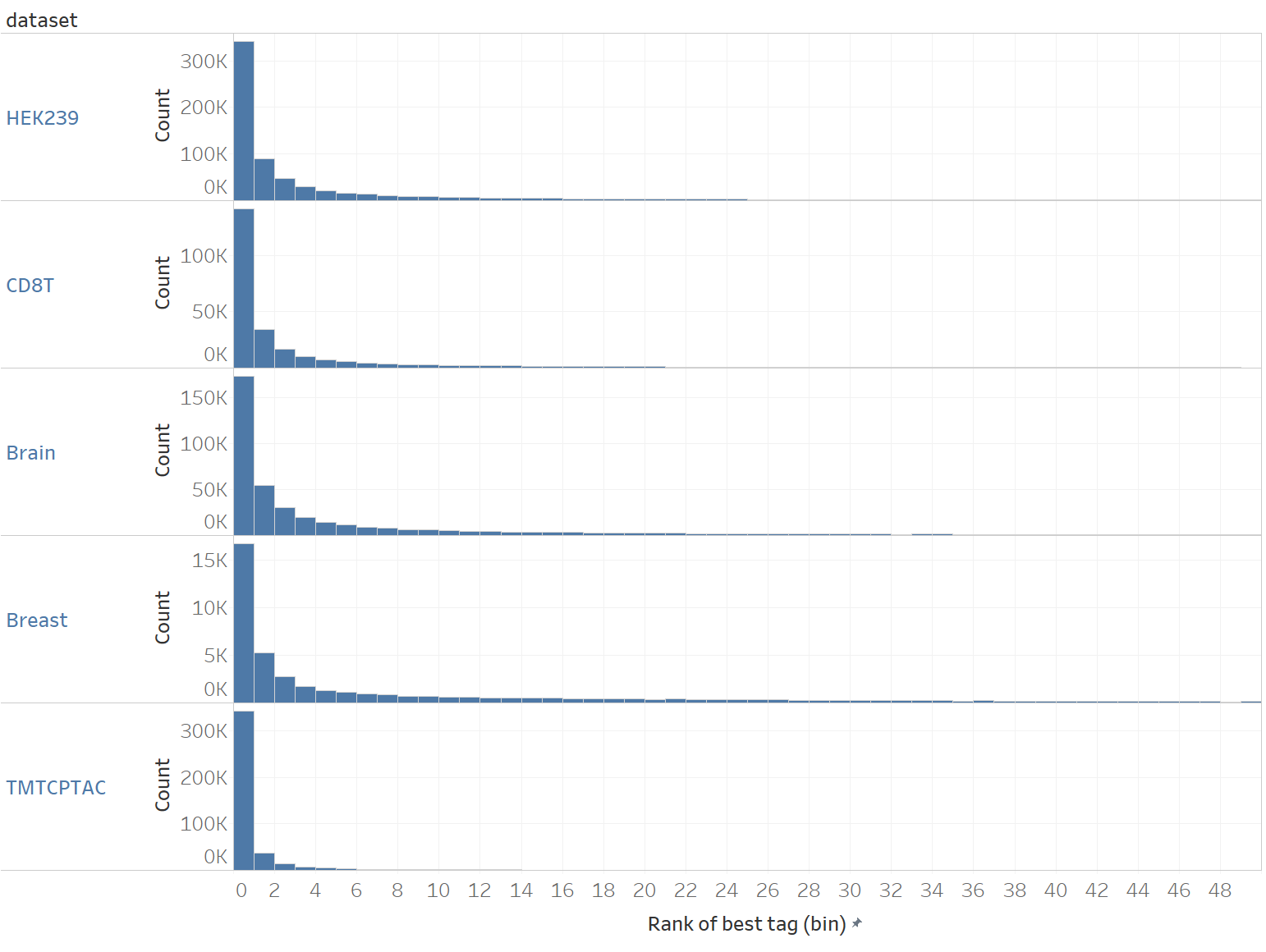

**Supplementary Fig. 2 | Distribution of tag ranks for PSMs identified in open search.** For each PSM, the rank of the highest ranked prefix or suffix tag is recorded as the rank for that PSM (x-axis).

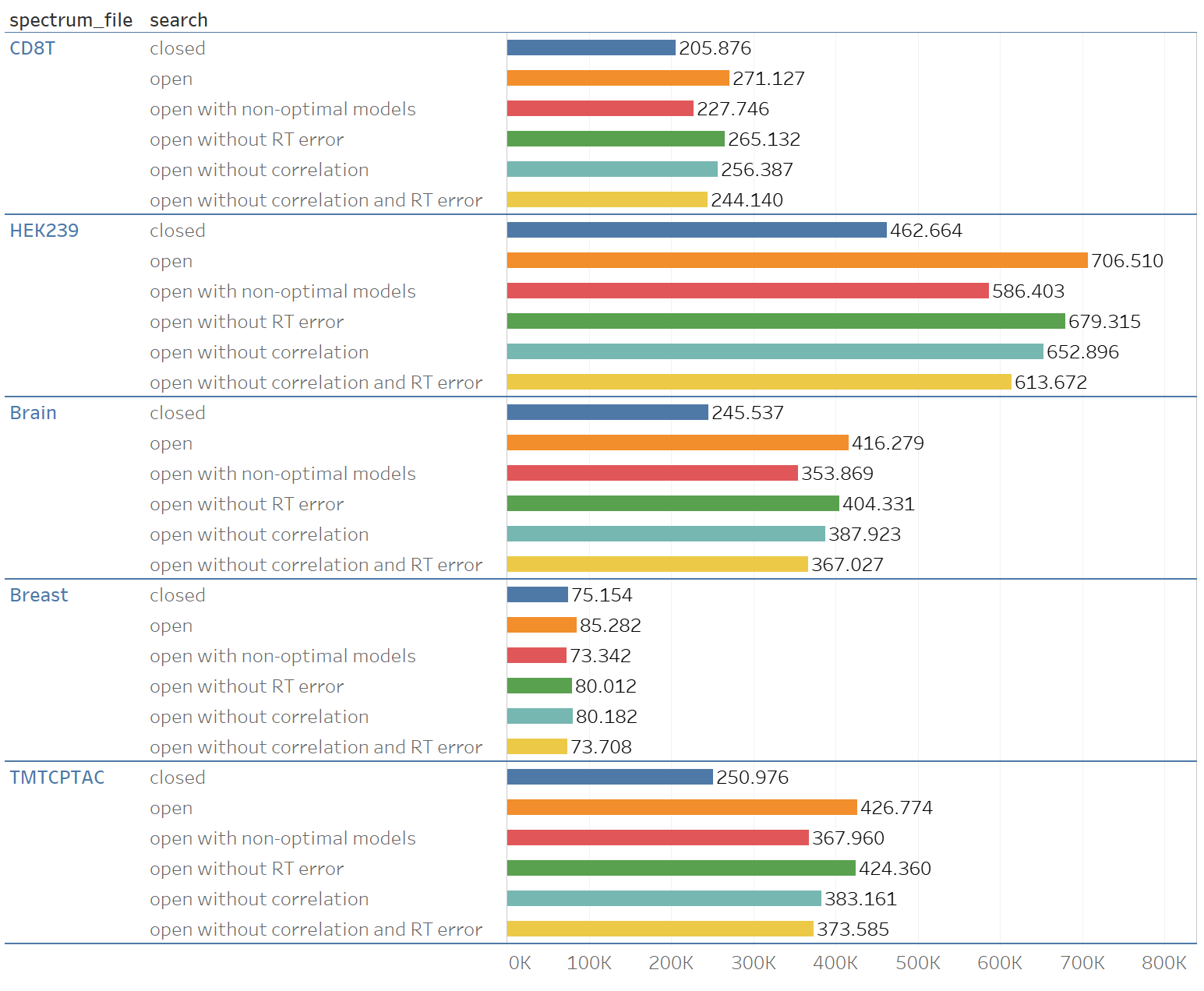

**Supplementary Fig. 3 | Number of identified PSMs (x-axis) for the five evaluation datasets using different search approaches.** First, a closed search (dark blue). Second, a standard open search (orange). Third, an open search using non-optimal models (red), i.e. for the TMTCPTAC dataset, models (tag-models, MS2PIP and DeepLC) trained on unlabeled HCD were used, while for all other datasets models trained on TMT labeled data were used. Fourth, an open search without using the RT-pred-error feature (green). Fifth, an open search without using the intensity-correlation feature and biased scoring function (light blue). Sixth, an open search without intensity-correlation and RT-pred-error (yellow).

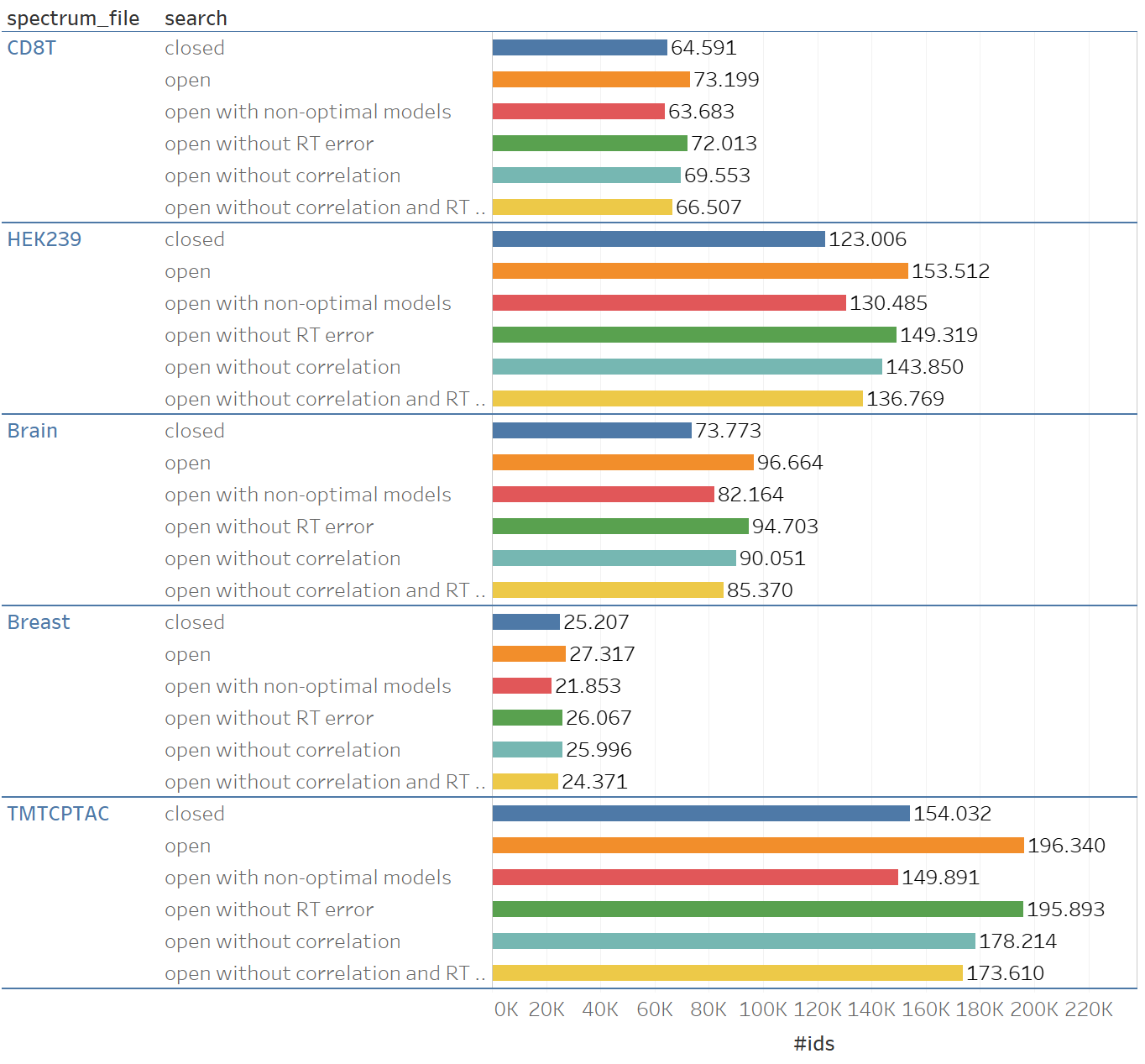

**Supplementary Fig. 4 | Number of identified peptides (x-axis) for the five evaluation datasets using different search approaches.** First, a closed search (dark blue). Second, a standard open search (orange). Third, an open search using non-optimal models (red), i.e. for the TMTCPTAC dataset, models (tag-models, MS2PIP and DeepLC) trained on unlabeled HCD were used, while for all other datasets models trained on TMT labeled data were used. Fourth, an open search without using the RT-pred-error feature (green). Fifth, an open search without using the intensity-correlation feature and biased scoring function (light blue). Sixth, an open search without intensity-correlation and RT-pred-error (yellow).

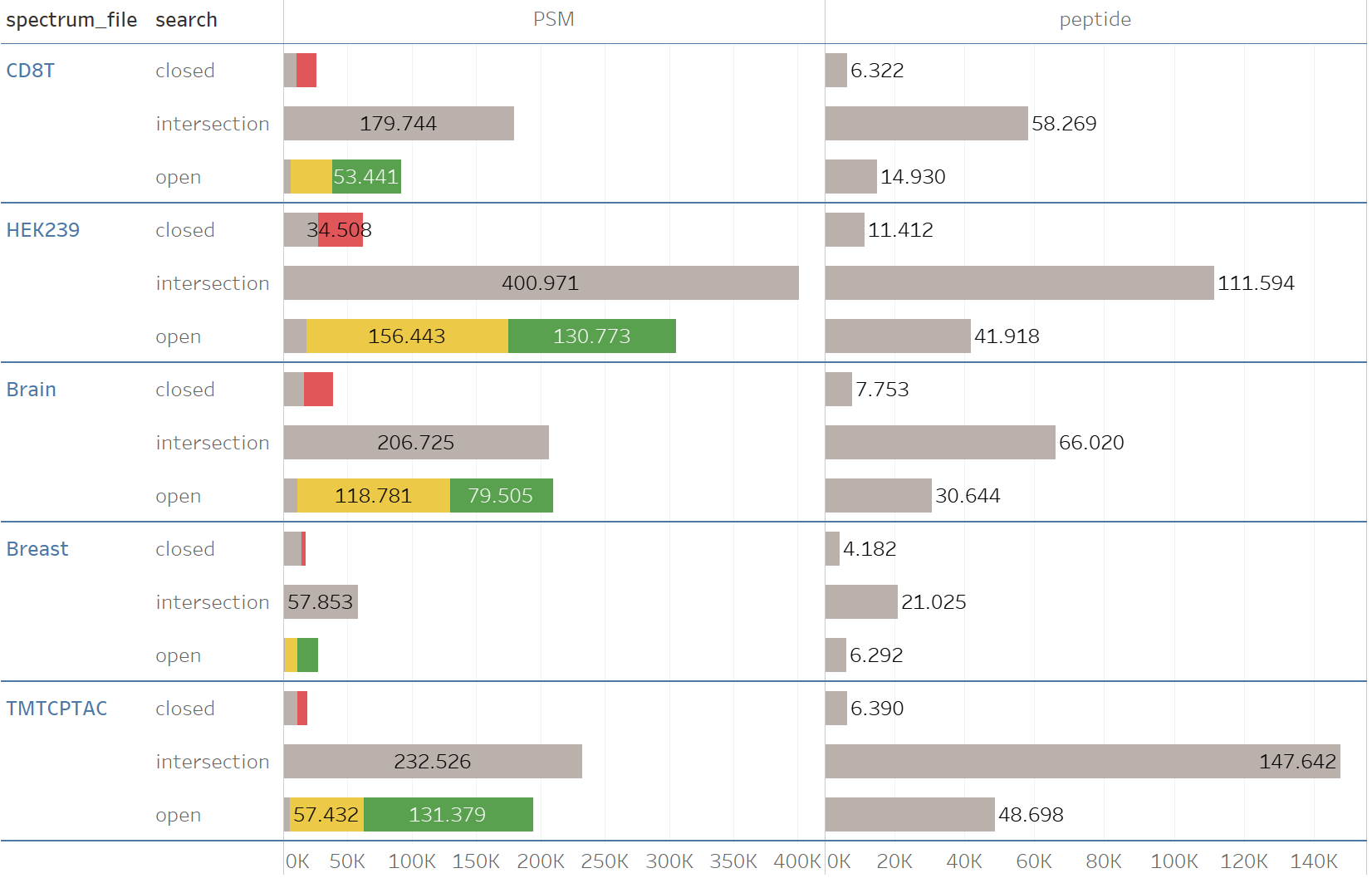

**Supplementary Fig. 5 | PSM and peptide identification overlap between a closed and open search.** For the closed searches, spectra with overruled identifications are shown in red. For open searches, matches with an unexpected modification are shown in green, while wide error matches are shown in yellow.

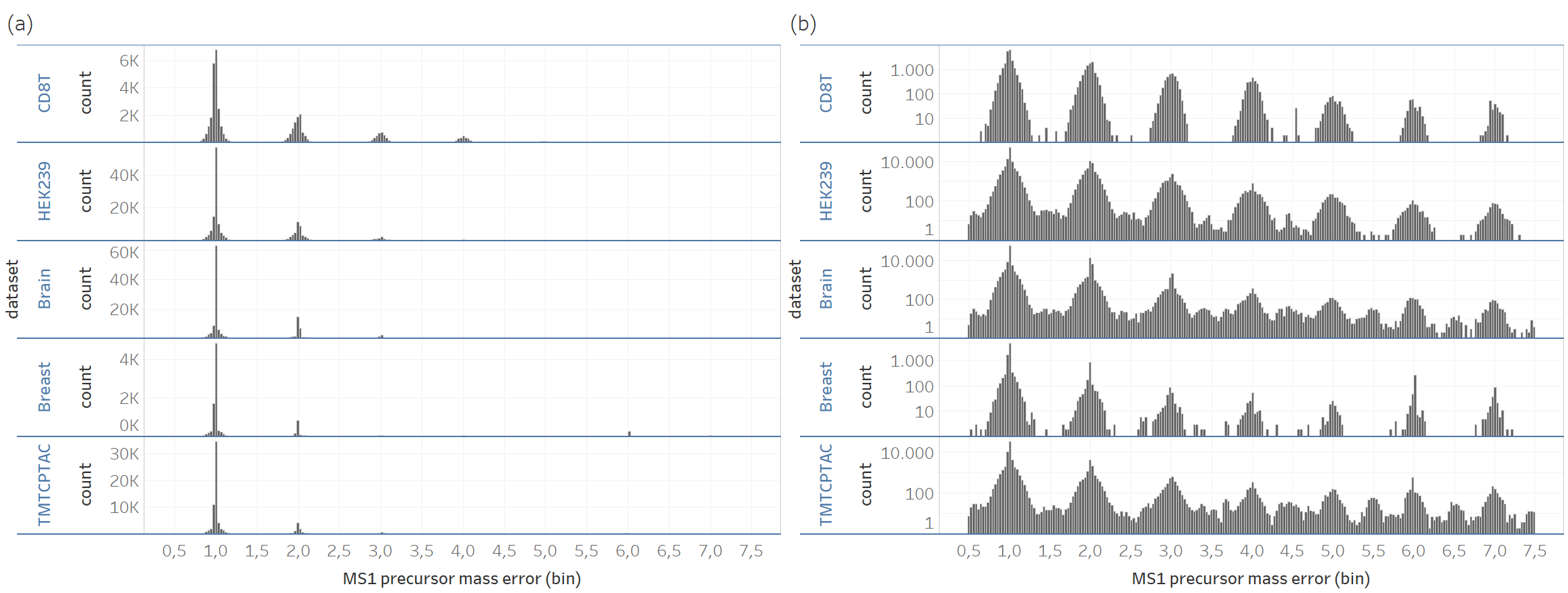

**Supplementary Fig. 6 |MS1 precursor error for wide error matches.** In **(a)** the MS1 precursor error distribution for PSM identifications with error larger than 0.5 Da is shown. In **(b)** the same information is shown as in (a) but with y-axis in log_10_-scale.

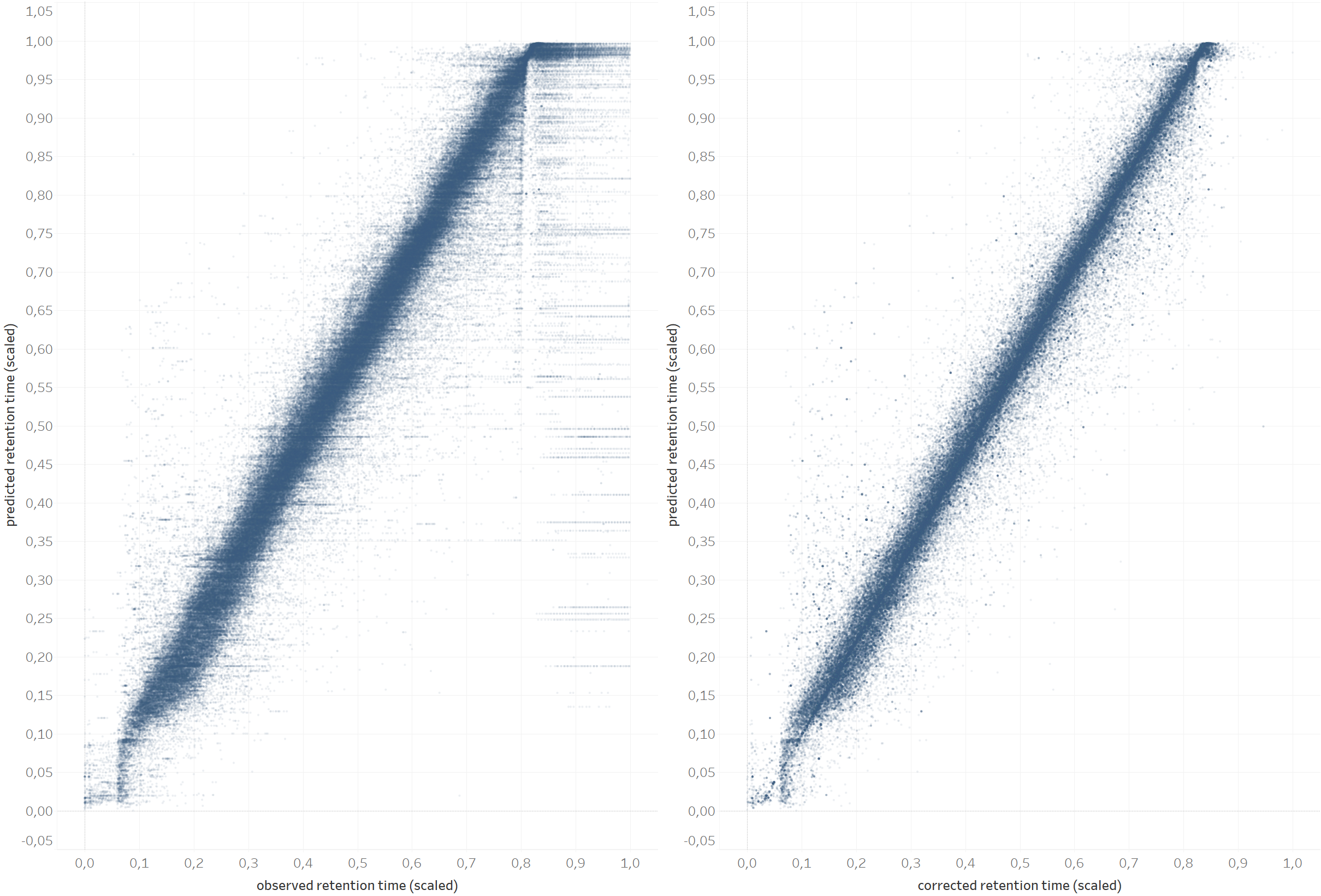

**Supplementary Fig. 7 | The effect of grouping peptidoforms to compute the corrected observed retention time.** Both plots show the observed versus predicted retention time for PSMs identified in an open ionbot search for CD8T. In **(a)** the true observed retention time for each PSM is plotted. In **(b)** the corrected observed retention time is plotted for the same dataset (CD8T). Retention times are normalized by dividing by the maximum observed retention time.

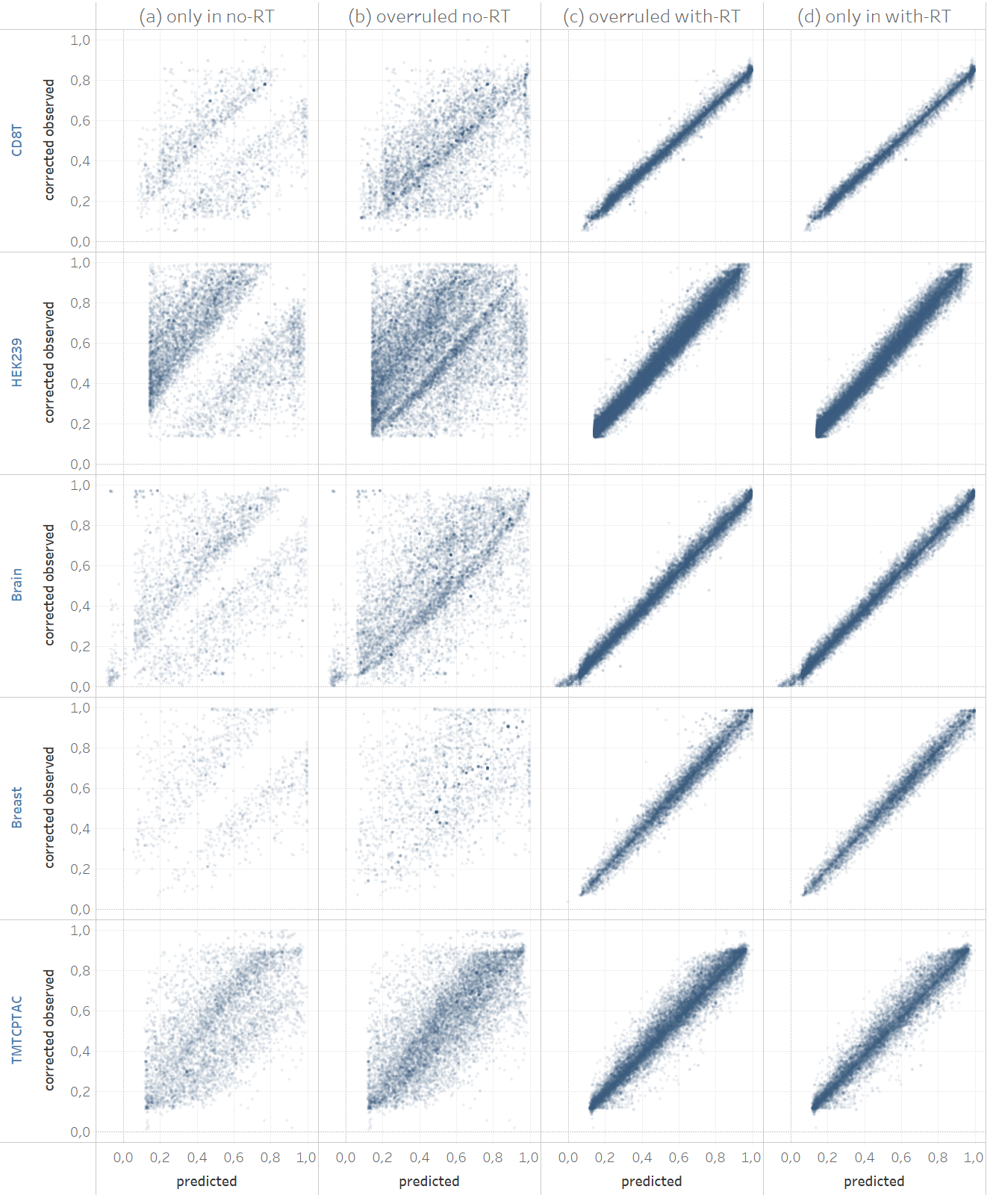

**Supplementary Fig. 8 | Corrected observed versus predicted retention times for PSMs identified in two ionbot open searches;** one does not the RT-pred-error feature in the PSM scoring function (no-RT), while the other performs a normal open search that does use this feature (with-RT). In **(a)** all matches identified in the no-RT search only are shown. In **(b)** all matches identified in the no-RT search that were overruled by another match in the with-RT search are shown. In **(c)** these overruled matches for the with-RT search are shown. In **(d)** all matches identified in the with-RT search only are shown.

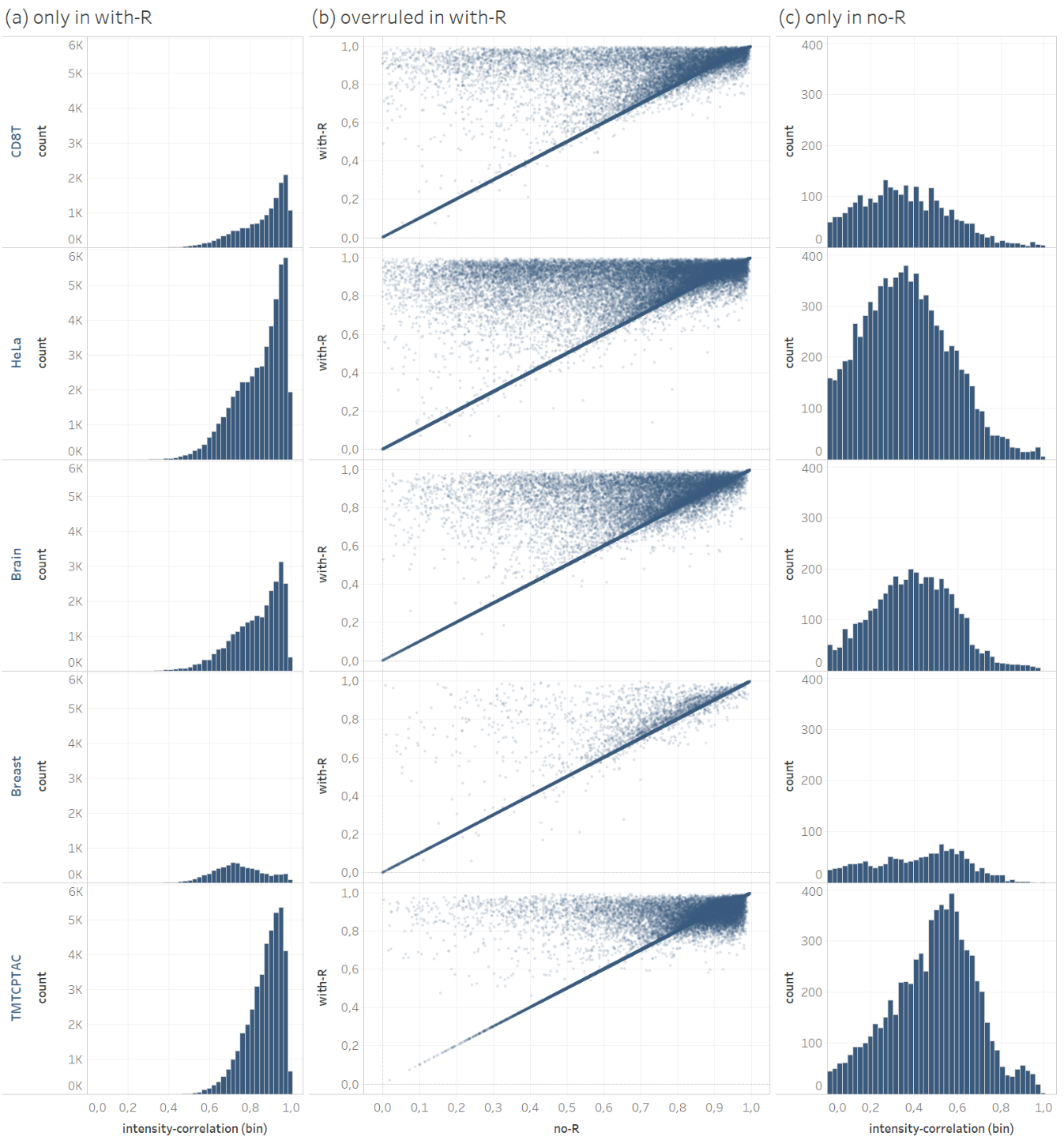

**Supplementary Fig. 9 | Comparing peak intensity pattern correlations for PSMs identified in two ionbot open searches;** one does not the intensity-correlation feature in the PSM scoring function or as a biased scoring function (no-R), while the other performs a normal open search that does use this feature (with-R). In **(a)** all matches identified in the no-R search only are shown. In **(b)** all PSMs identified in the no-R search that were overruled by another match in the with-R search are plotted. For each such PSM, the intensity-correlation of the match in the no-R search (x-axis) is plotted against the intensity-correlation of the match in the R search (y-axis). In **(c)** all matches identified in the with-R search only are shown.

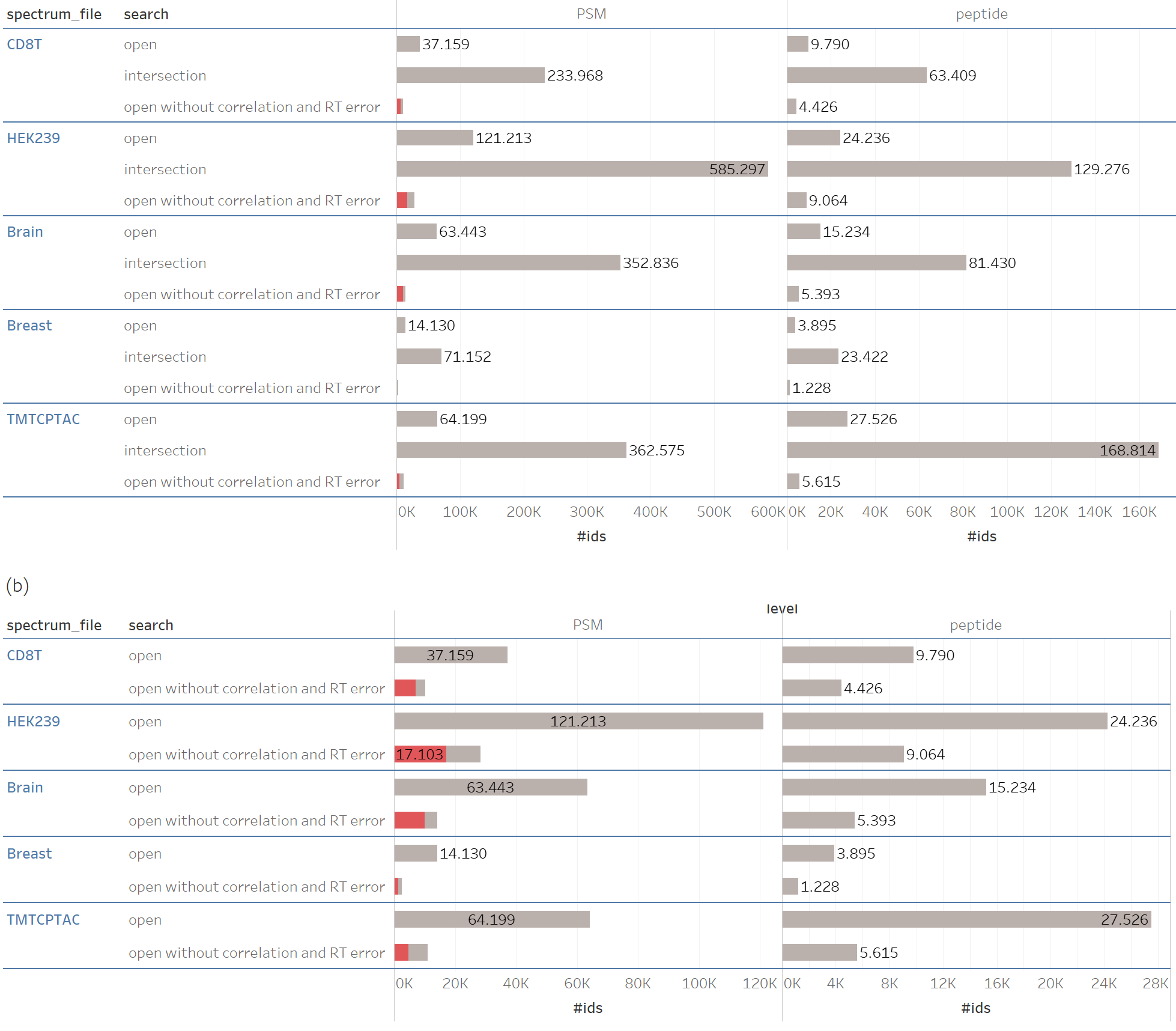

**Supplementary Fig. 10 | Comparing a normal ionbot open search with an open search that does not use the RT-pred-error and the intensity-correlation in the PSM scoring function or as a biased PSM scoring function.** Matches overruled in the normal open search are shown in red. The first column shows identification overlap at PSM level, while the second column shows identification overlap at the unique peptide level. The bars in **(a)** show the same counts as in **(b)**, but also shows the number of matches that were the same in both searches (intersection).

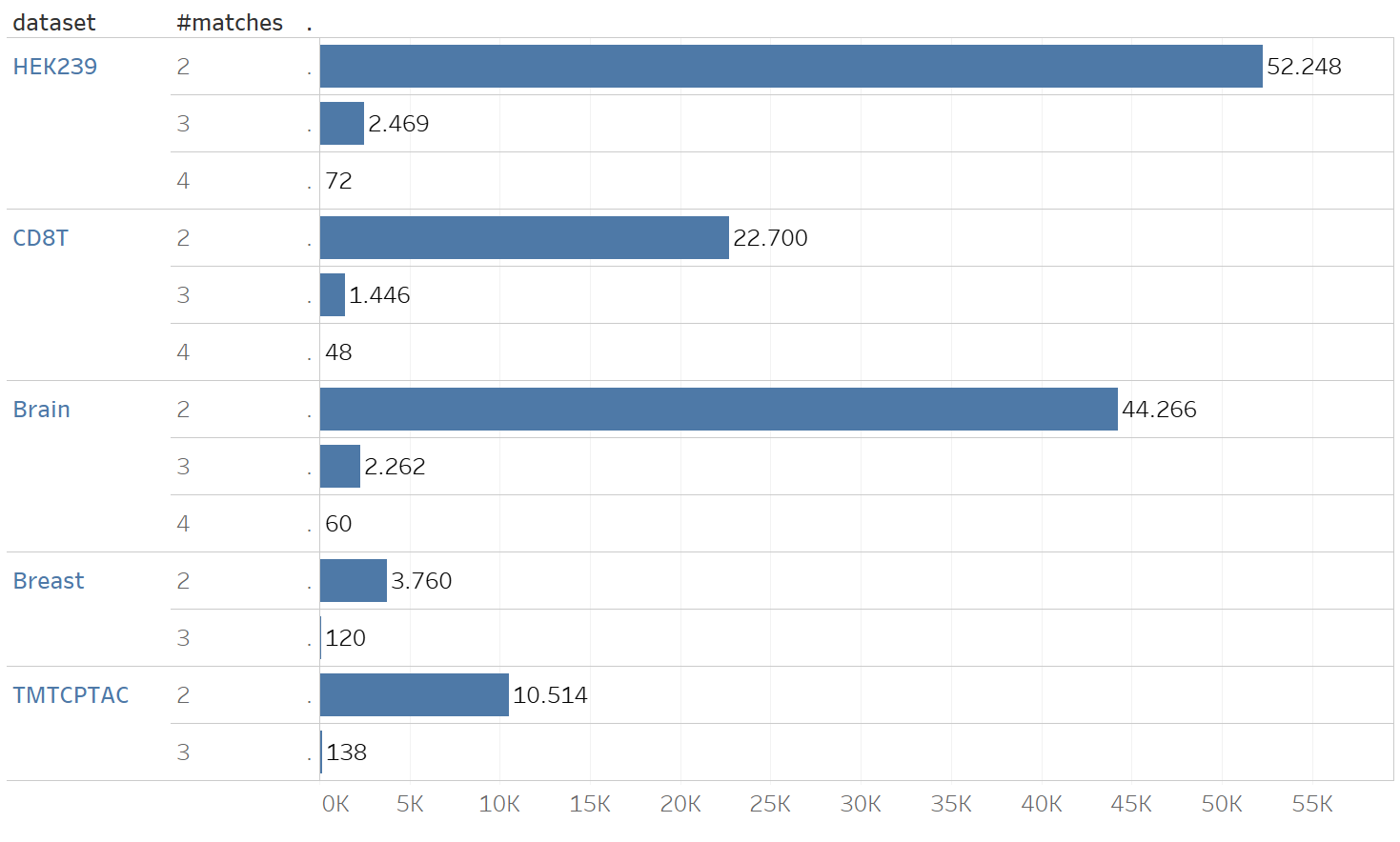

**Supplementary Fig 11 | For each of the five evaluation datasets the bars count the number of spectra with 2 or more (#matches) co-eluting matches.** Spectra with a single match are not shown.

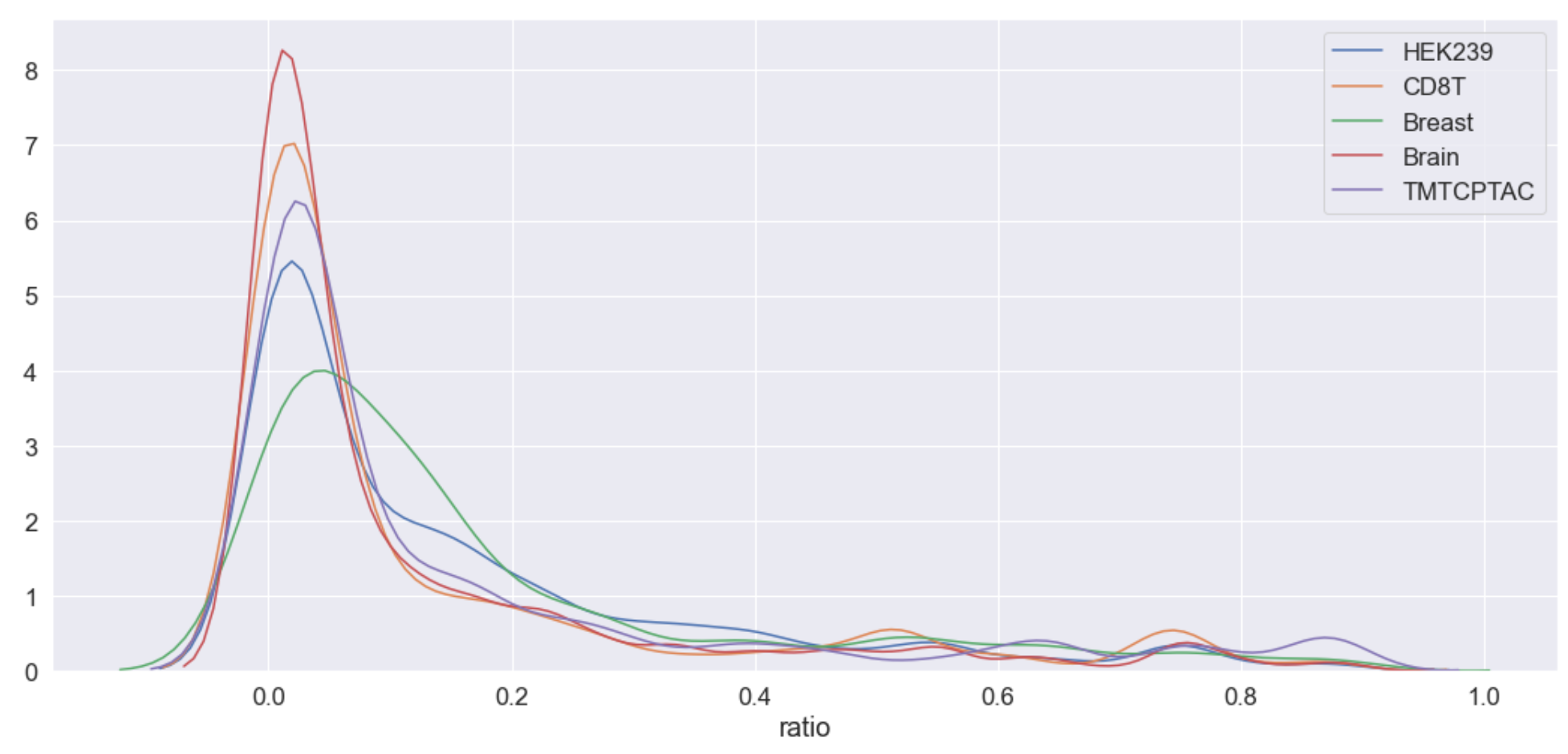

**Supplementary Fig. 12 | The b- and y-ion overlap between co-eluting matches.** The ratio (x-axis) is computed as the intersection of the b- and y-ion peak masses over the union of these masses and plotted as a density plot. All pairwise combinations of co-eluting matches are shown.. Both histogram, with bin size equal to one, and density plot are shown.

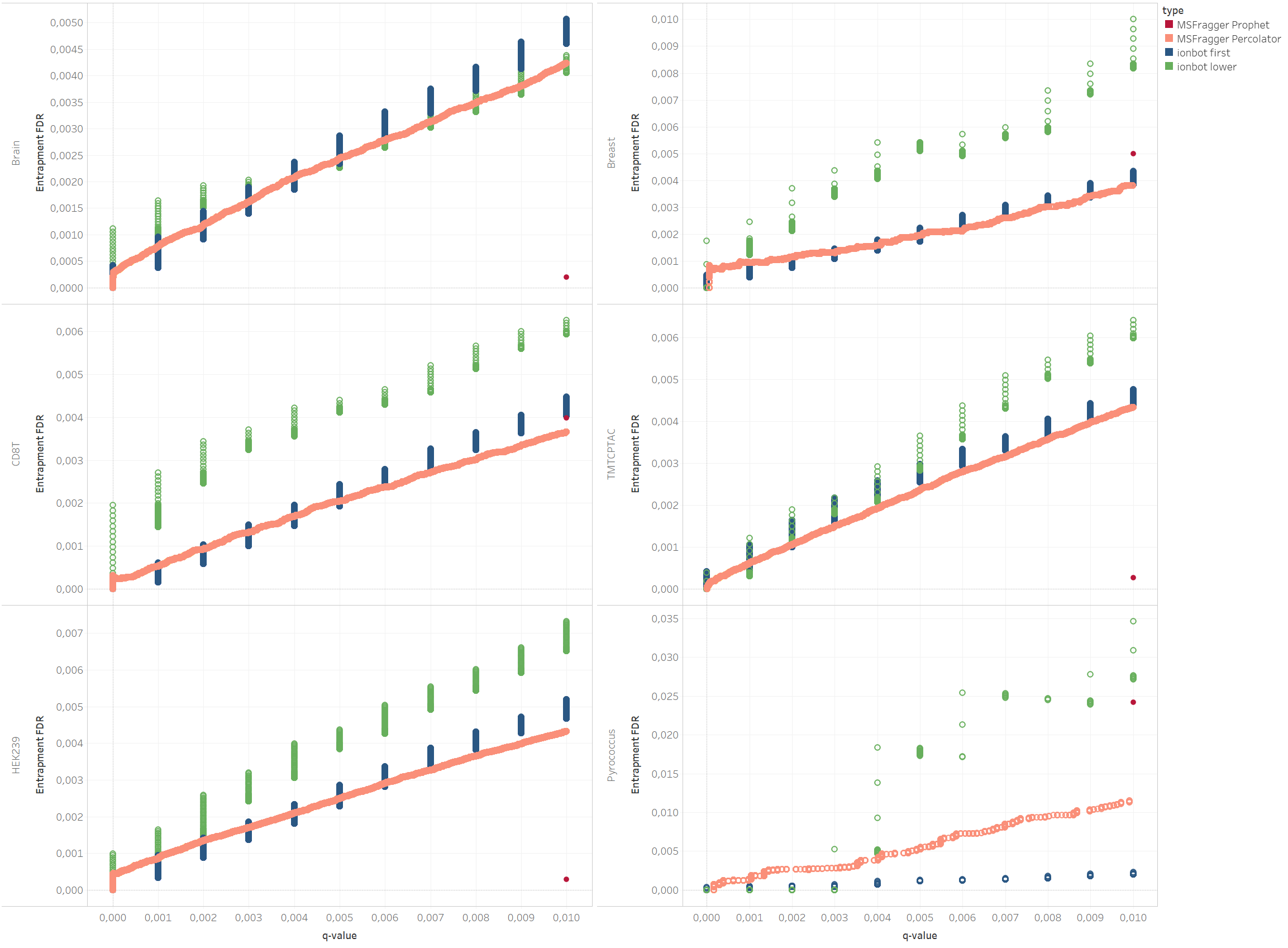

**Supplementary Fig. 13 | The entrapment FDR (y-axis) plotted against the PSM q-values (x-axis) for the five evaluation datasets and the *Pyrococcus furiosus* dataset.** For MSFragger results are shown with PeptideProphet (MSFragger Prophet) and Percolator (MSFragger Percolator) post-processing. For ionbot the entrapment FDR is computed separately for the first (ionbot first) and the lower (ionbot lower) ranked matched. For MSFragger Prophet no statics are returned my MSFragger, resulting in just one FDR datapoint (in red) at q-value = 0.01.

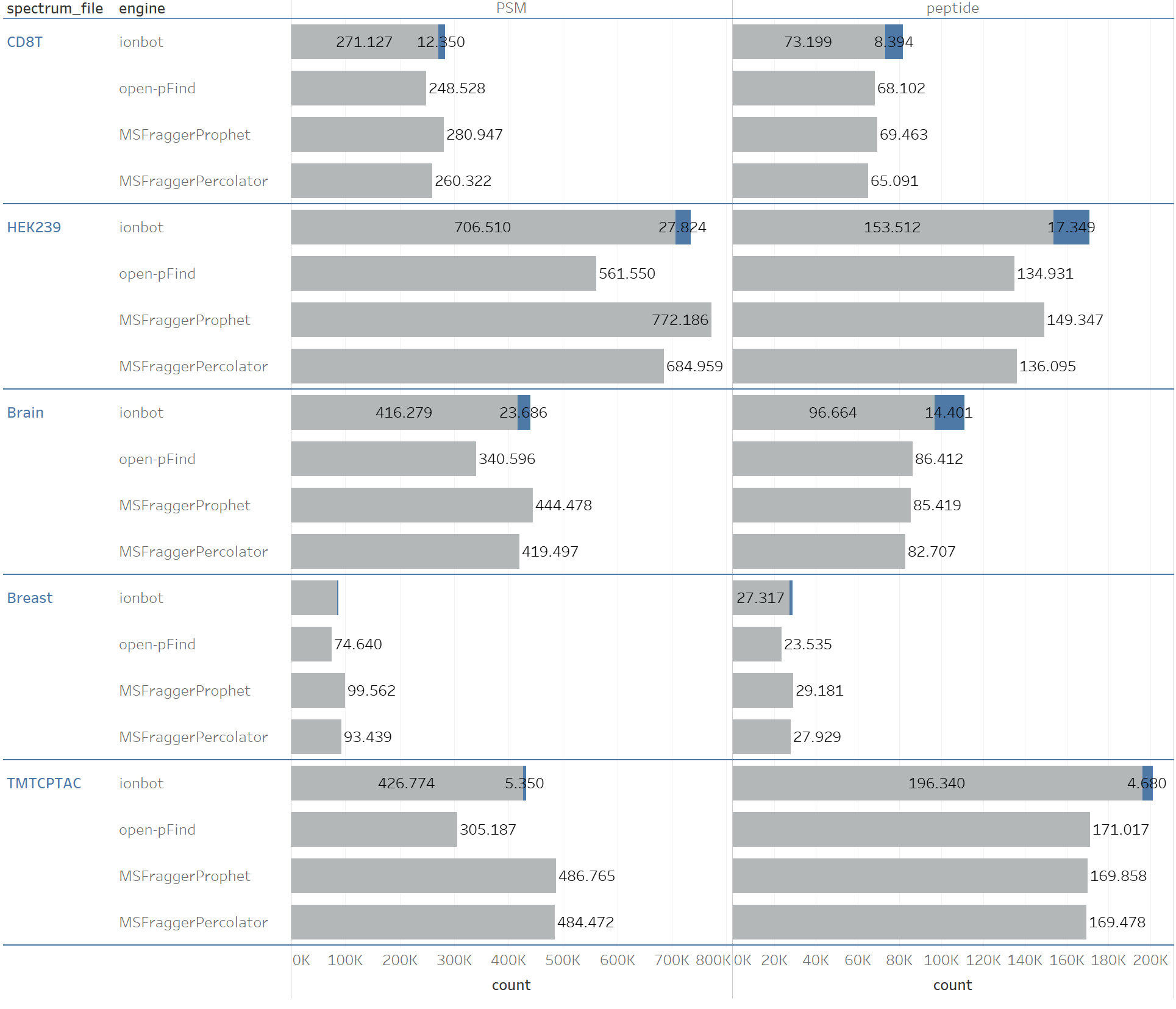

**Supplementary Fig. 14 | The number of PSM (first column) and unique peptide (second column) identifications for each search engine.** For ionbot, co-eluting lower ranked matches are shown as the blue sections in the bars. The first column shows PSM identifications, the second column shows uniquely identified peptide sequences.

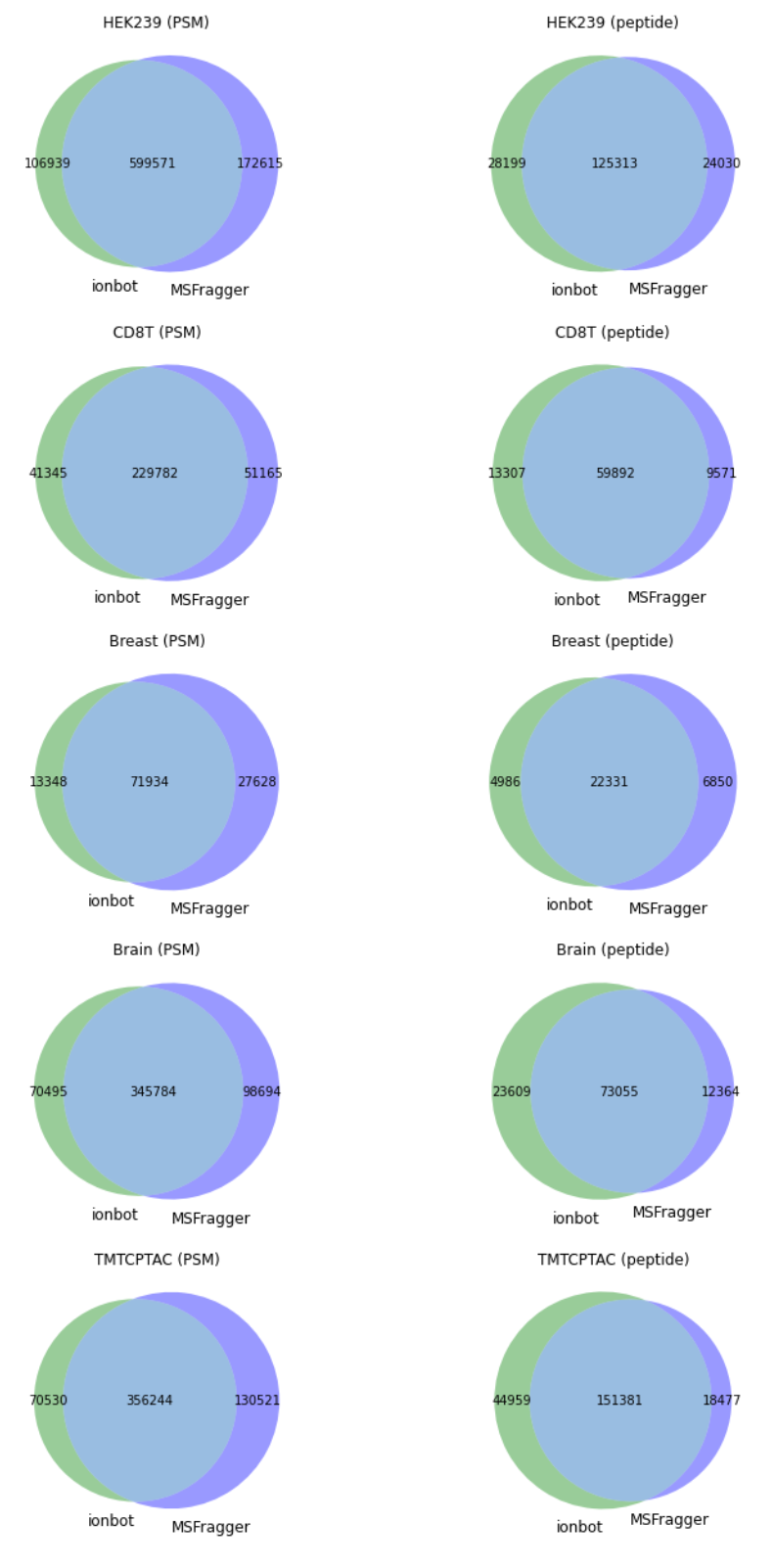

**Supplementary Fig. 15 | The identification overlap between three open modification identification engines at PSM level (first column), and at unique peptide sequences level (second column).**

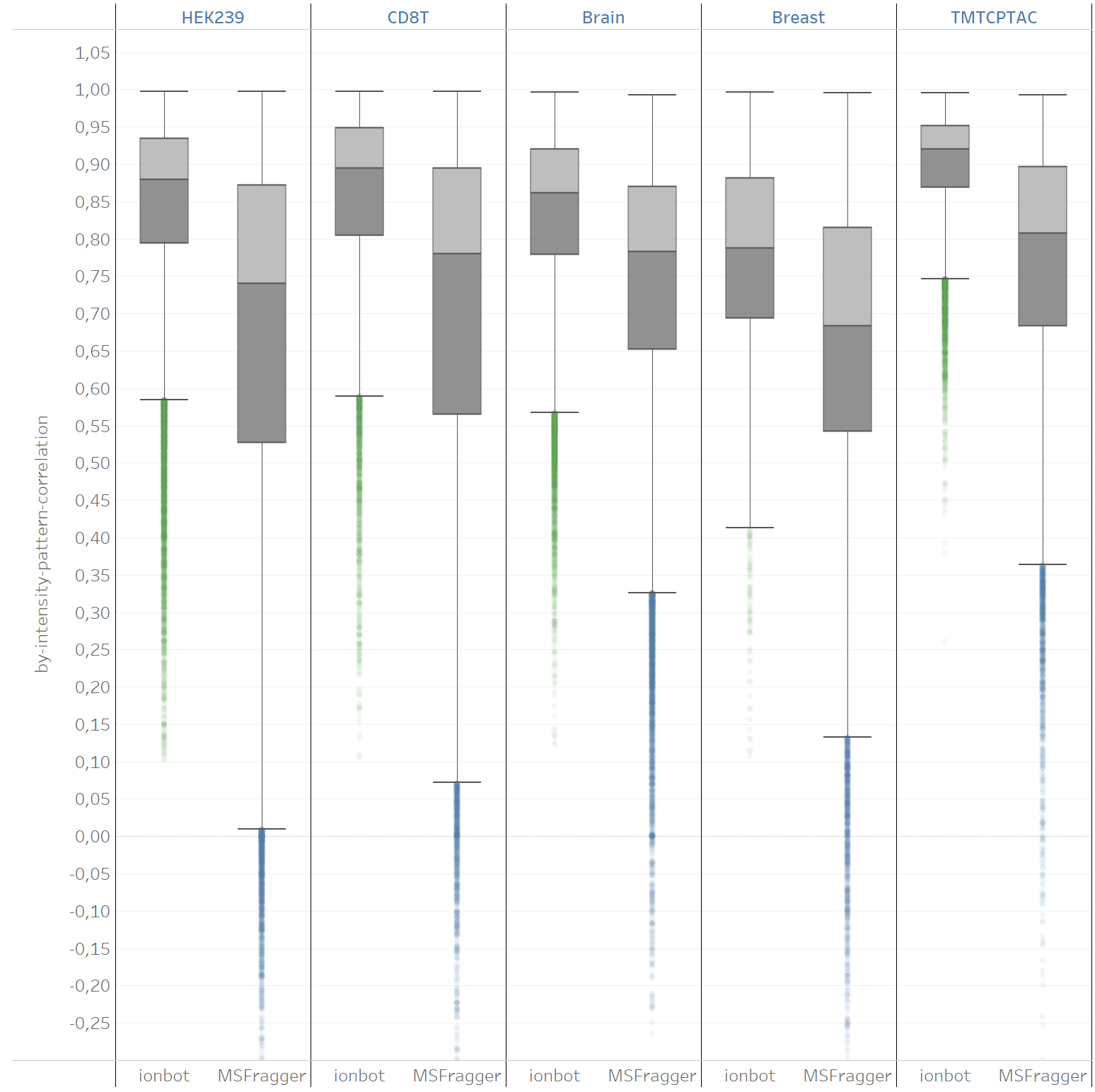

**Supplementary Fig. 16 | The value of the intensity-correlation feature (y-axis) for matches unique to ionbot or MSFragger.** Only matches that don’t contain an unexpected modification as shown.

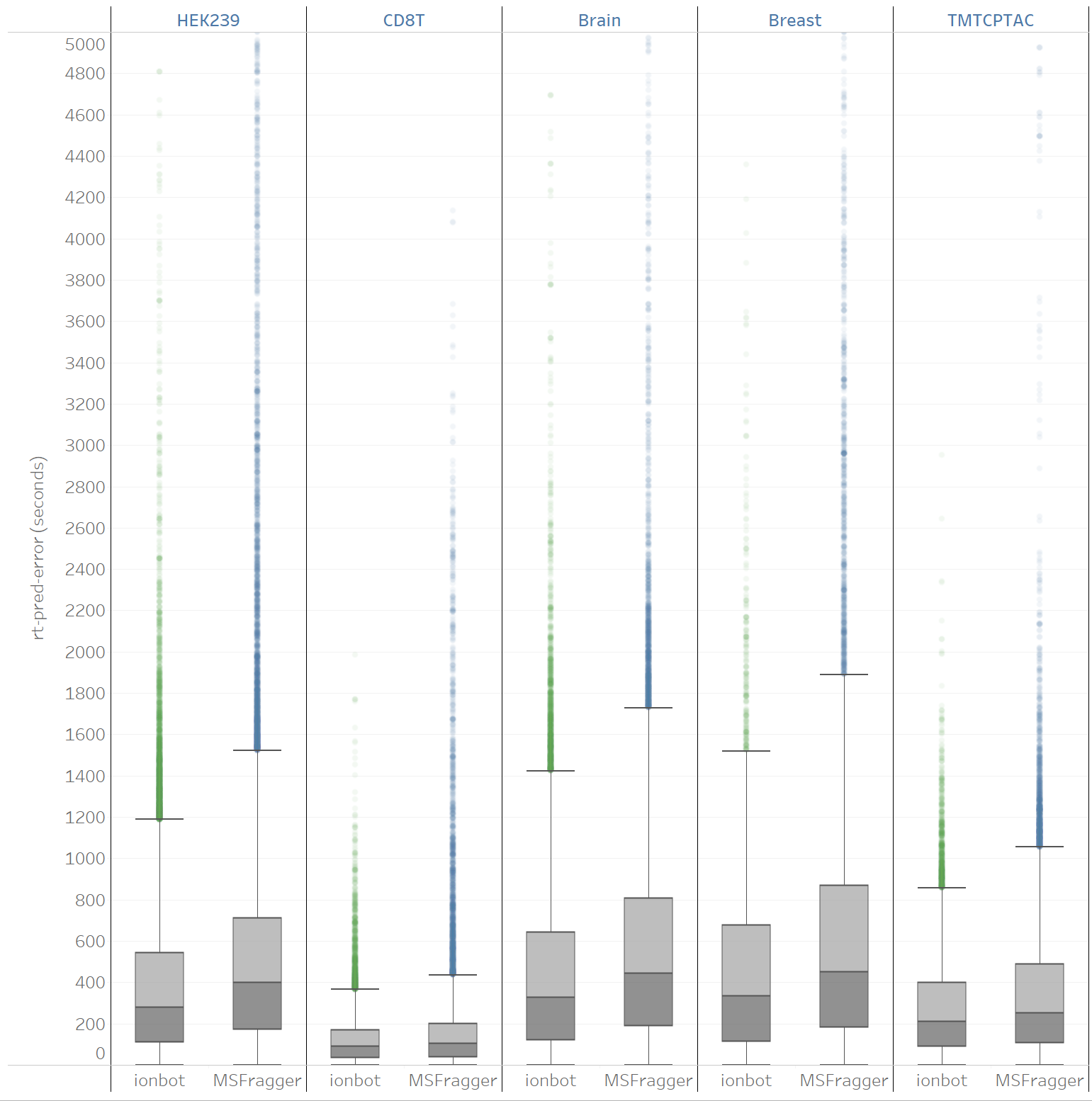

**Supplementary Fig. 17 | The value of the RT-pred-error feature for matches unique to ionbot or MSFragger.** Only matches that don’t contain an unexpected modification as shown.
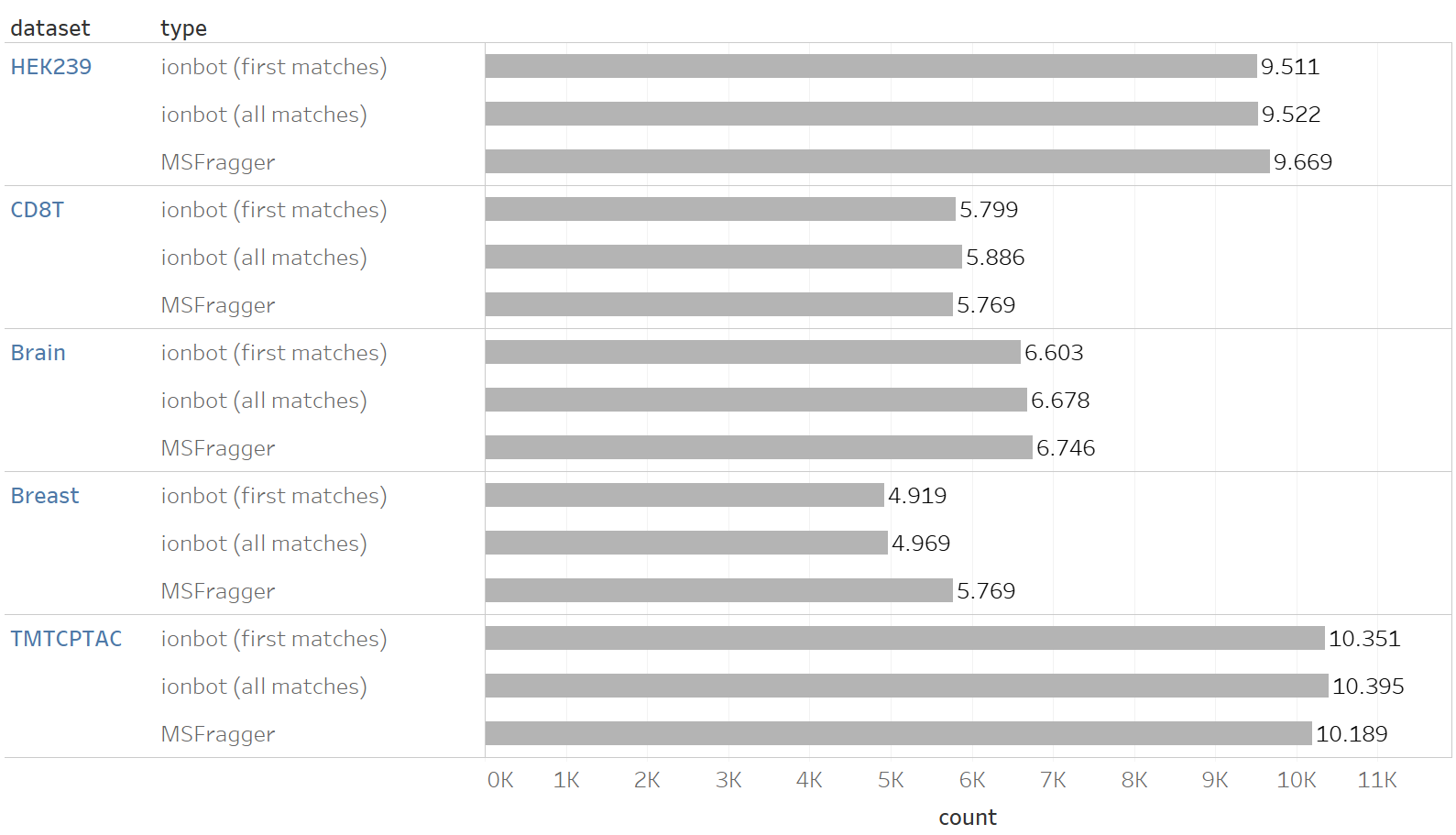

**Supplementary Fig. 18 | The number of protein group identifications for ionbot and MSFragger.** The results for ionbot are split into proteins identified using the first ranked PSMs only (first matches), and proteins identified with the co-eluting matches as well (all matches). For MSFragger, ProteinProphet was used for protein inference.
